# Compensatory mutation can drive gene regulatory network evolution

**DOI:** 10.1101/2019.12.18.881276

**Authors:** Yifei Wang, Marios Richards, Steve Dorus, Nicholas K. Priest, Joanna J. Bryson

## Abstract

Gene regulatory networks underlie every aspect of life; better understanding their assembly would better our understanding of evolution more generally. For example, evolutionary theory typically assumed that low-fitness intermediary pathways are not a significant factor in evolution, yet there is substantial empirical evidence of compensatory mutation. Here we revise theoretical assumptions to explore the possibility that compensatory mutation may drive rapid evolutionary recovery. Using a well-established *in silico* model of gene regulatory networks, we show that assuming only that deleterious mutations are not fatal, compensatory mutation is surprisingly frequent. Further, we find that it entails biases that drive the evolution of regulatory pathways. In our simulations, we find compensatory mutation to be common during periods of relaxed selection, with 8-15% of degraded networks having regulatory function restored by a single randomly-generated additional mutation. Though this process reduces average robustness, proportionally higher robustness is found in networks where compensatory mutations occur close to the deleterious mutation site, or where the compensatory mutation results in a large regulatory effect size. This location- and size-specific robustness systematically biases which networks are purged by selection for network stability, producing emergent changes to the population of regulatory networks. We show that over time, large-effect and co-located mutations accumulate, assuming only that episodes of relaxed selection occur, even very rarely. This accumulation results in an increase in regulatory complexity. Our findings help explain a process by which large-effect mutations structure complex regulatory networks, and may account for the speed and pervasiveness of observed occurrence of compensatory mutation, for example in the context of antibiotic resistance, which we discuss. If sustained by *in vitro* experiments, these results promise a significant breakthrough in the understanding of evolutionary and regulatory processes.

## 1 Introduction

Gene regulatory networks are the internal tools living organism employ to facilitate resilience to environmental variation. Unfortunately, to date their evolution is not well understood [Wilke and Adami, 2001, Wilke et al., 2003, Beerenwinkel et al., 2007, Lehner, 2011, Rokyta et al., 2011, Park and Lehner, 2013]. Prior theory building based on computational and mathematical models has shown that gene regulatory networks (GRNs) can evolve by adaptive responses to direct selection [Ciliberti et al., 2007, Crombach and Hogeweg, 2008, Romero and Arnold, 2009, Tsuda and Kawata, 2010, Olson-Manning et al., 2012, Cotterell and Sharpe, 2013], and by random genetic drift when sampling biases shift the distribution of networks [cf. Wagner and Wright, 2007, Lynch et al., 2016].

In contrast, the role of compensatory mutation in GRN evolution has been given little consideration. In most modelling, regulatory networks rendered unfunctional (unstable) by deleterious mutation have been assumed to be immediately purged by direct selection, precluding the possibility of an additional mutation which might restore fitness. Yet in nature, phenotypic selection is known to be episodic, with periods of strong and weak selection [Siepielski et al., 2009a]. This indicates there may be sufficient time for evolutionary rescue. Further, if compensatory mutation happens frequently enough, it could play a major role in GRN evolution. Compensatory mutation has been observed to occur surprisingly frequently in some laboratory contexts under high selective pressure, particularly with respect to antibiotic resistance [Dunai et al., 2019, Moura de Sousa et al., 2017], see Remigi et al. [2019] for a recent review. This indicates that compensatory mutation should receive more theoretical attention.

Compensatory mutation could contribute to GRN evolution as an emergent consequence of biases that occur in the processes of mutation and selection. Although originally considered an important source of innovation and diversity, mutations are now thought generally to be in the vast majority of cases at least mildly deleterious, decreasing individual fitness. However, not all mutations are deleterious or have the same detrimental effects on all individuals. There are occasionally beneficial mutations, including compensatory mutations that recover fitness after deleterious mutations [Kulathinal et al., 2004, Piskol and Stephan, 2008, Covert et al., 2013]. Such mutations could contribute to gene pathway evolution [Kimura, 1985, Moore et al., 2000, Levin et al., 2000, Choi et al., 2005, Meer et al., 2010]. However, in contrast to adaptive and neutral evolution, little attention has been placed on evolutionary dynamics driven by genes that at least initially code lower-than-average reproductive success.

Compensatory mutation in regulatory networks may be far more frequent than we expect. Outside of the context of a regulatory network, theory tells us compensatory mutation is not likely to play an important role in evolution [Wright, 1931a,b, Stephan, 1996, Parsch et al., 1997, Whitlock and Otto, 1999, Whitlock et al., 2003, Zhang and Watson, 2009]. This is because the frequency of deleterious mutation is low and the frequency at which a new mutation compensates for the previous deleterious mutation is expected to be even lower. Furthermore, if the compensatory mutation restores fitness, then its probability of fixation in the population might be assumed to be the same as any allele under drift, the inverse of twice the effective population size [Wright, 1931a, Charlesworth, 2009]. This analysis may be incorrect though, given the coupling to the original deleterious mutation, particularly if the ‘deleterious’ mutation provides a specialist ability to an organism despite overall lowering its adaptive value. This is the reported case in antibiotic resistance, where initial mutations confer resistance but create other metabolic costs. Some proportion of bacteria evolve compensatory mutations rather than reverting to initial genotype [Dunai et al., 2019], and some even seem to be conferred with net adaptive advantage after the event of additional mutations [Moura de Sousa et al., 2017]. Further, mutations do not only happen in independently-acting genes, but also in genetic networks where there are many sites of complex interactions that could be mutated. If a deleterious mutation occurs at a locus that is not presently subjected to strong selective pressure, then as long as a compensatory mutation occurs before the lineage is driven to extinction, it may restore the lineage’s fitness. Thus, understanding the frequency and nature of compensatory mutations is of substantial importance to understanding their impact on pathway evolution.

Logically, we can expect relaxed selection to be critical to the frequency of compensatory mutation. Empirical evidence indicates that when selection against deleterious mutation is relaxed, the frequency of compensatory mutation in organisms carrying deleterious mutations is surprisingly high [Maisnier-Patin et al., 2002, Gifford and MacLean, 2013]. Several recent empirical studies have suggested that various sorts of relaxed selection facilitate compensatory mutation [Sloan et al., 2014, Moura de Sousa et al., 2017, Dunai et al., 2019]. Compensatory mutations might then be expected to play a key role in the formation of GRN. The frequency at which deleterious mutations incapacitate gene regulatory pathways is likely to be substantially higher than that for an independently acting gene, because there will inevitably be many more possible sites to mutate. We do not know the frequency at which mutations in incapacitated networks can compensate for previous deleterious mutations. But because mutation, by definition, occurs in networks that were previously functional, it seems logical that there could be a wide range of mutational sites and magnitudes that might restore the function of a network. If the frequency of compensatory mutation is high and persistent enough over time, then there is a high probability that some compensatory mutations will be maintained, even if solely by drift. If there is something special about the sorts of genes likely to produce compensation, then we might also expect this process to generate biases, both with respect to where mutations occur and any other characteristic that might engender higher robustness. Variation in the speed with which poorly functioning genotypes are removed by purifying natural selection could ultimately have substantial impacts on the genetic attributes of the population. Similarly, properties associated with compensatory mutations may accumulate over time. The basic logic of this argument is that, facilitated by periods of relaxed selection, compensatory mutation not only allows evolution to proceed, but results in it proceeding *differently*. GRN may accumulate specific features as a consequence of compensatory mutation.

The gene regulatory network paradigm is an excellent system is which to test this question. Most importantly, it explicitly incorporates genetic interactions in an evolutionary framework. Simulation allows us to generate thousands upon thousands of networks of different sizes and connectivities, which we could not do with *in vivo* approaches, and makes it relatively simple to identify, track and understand the properties of all of the compensatory mutations within those networks. Many previous computational studies have focused on the evolution of gene regulatory networks under constant selection [Azevedo et al., 2006, Ciliberti et al., 2007, Crombach and Hogeweg, 2008, Tsuda and Kawata, 2010, Cotterell and Sharpe, 2013]. However, constant selection necessarily constrains pathway evolution because it removes the low-fitness individuals who carry incapacitated gene networks. This in turn eliminates the potentially significant mechanism of compensatory mutation. Compensatory mutation is impossible under one of the dominant modelling frameworks, where unstable networks — networks whose phenotype never reach an equilibrium state — are always labelled as ‘unviable’ and therefore never subjected to further rounds of mutation [Wagner, 1996, Siegal and Bergman, 2002, Azevedo et al., 2006, Lohaus et al., 2010]. If we instead allow unstable networks to stay in the population when selection for network stability is relaxed, compensatory mutation is possible and able to allow lineages access to a greater variety of evolutionary pathways.

In our experiments, we adapted the experimental paradigm for one of the dominant models of GRN evolution, to consider deleterious mutations to only compromise, not destroy, a lineage. This allows us to examine the prevalence and impacts of compensatory mutation. We find that compensatory mutations are both surprisingly common, and that their frequency is relatively invariant to the scale of the network. We also find that compensatory mutations exhibit biases both in effect size, and in location with respect to the deleterious mutation. The accumulation of these emergent biases increases regulatory complexity in GRN over generations. As we discuss, these observations are all congruent with empirical observations, indicating we may have established a useful theoretical advance in the understanding of compensatory mutation and of GRN.

## Results

### A network model for compensatory evolution

We present a model to study compensatory mutation, using a process well-established in the literature (see more details in Methods). As with published precedent on gene regulatory network (GRN) evolution, the model generates an initial population of stable regulatory networks and networks made unfunctional (unstable) by deleterious mutation [Wagner, 1996, Siegal and Bergman, 2002, Azevedo et al., 2006, Wang et al., 2015, Wang, 2019a,b]. Our innovation is that instead of assuming that unfunctional networks are removed immediately by persistent selection for network stability, we assume that they are part of a larger organism and only marginally reduce that organism’s fitness. In this, we effectively hypothesise that organisms carry a deleterious mutation (DM) load, analogous to parasite load. We compute the consequences of additional rounds of mutation on the stability of the network, during periods labelled as bouts of relaxed selection (Fig. 1).

**Figure 1:**
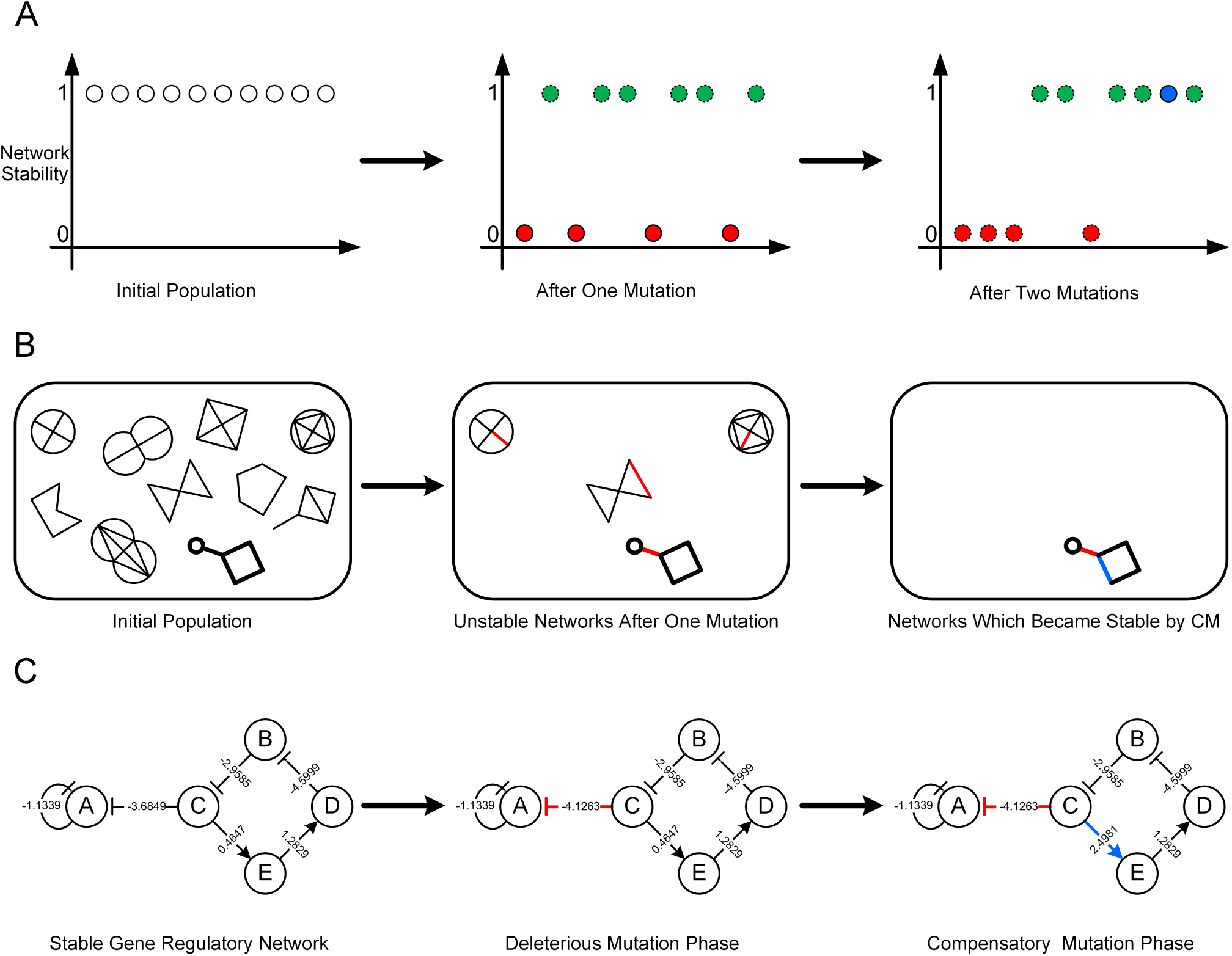
Overview across time of the computational model for exploring characteristics of compensatory mutation. (**A**) Stability (approximates fitness contribution) of gene regulatory networks in a population. Note that dashed circles are networks no longer considered for the study. (**B**) The population pool of gene regulatory networks. Note that a red edge indicates a deleterious mutation and a blue edge a compensatory mutation. (**C**) Detailed view of a single network.

Though the criteria for network stability we employ is similar to that used in previous models, our fitness function—the relationship between genotype and fitness—is different. Specifically, we assume that networks are either functional (high fitness, equivalent to fitness 1 in previous models) or unfunctional (low fitness, replacing fitness 0 in previous models). We then estimate the rate of compensatory mutation by calculating the proportion of regulatory networks which have stability restored by a single additional mutation, represented by the blue circle in Fig. 1A. Note that the green circle represents that the network has experienced a neutral mutation that does not affect network stability. Thus in Step 1 we generate a random population of GRNs. In Step 2 (Fig. 1B), each GRN has been mutated (red edge) and the resulting unstable networks have been collected for further testing. In Step 3, the unstable networks have undergone a second round of mutation, allowing the collection and analysis of any newly-stable networks. In this case, one network’s mutation has been compensatory (blue edge).

Fig. 1C shows an initially-stable gene network which contains five genes: *A*–*E*. Each edge is directed and indicates the strength (weight) of the influence on one gene of another. In the Deleterious Mutation Phase, a mutation occurs on 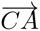 (red edge), which leads to the failure of maintaining network stability. In the Compensatory Mutation Phase, the compromised network is recovered by another round of mutation, with a mutation that proves compensatory (blue edge) occurring on 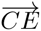. Note that there is no difference in the modelling process between generating a compensatory or deleterious mutation. Rather, random changes to the network are categorised based on their impact on network stability.

### Compensatory mutations are common and relatively scale-invariant

We first test whether compensatory mutation is common in the context of the synthetic GRNs. We find that, unlike deleterious mutation, the frequency of compensatory mutation is almost scale-invariant. From Fig. S1A and B, we can see that the stability and robustness in initial networks are quite different among varying sizes and levels of connectivity of gene regulatory networks. Which type of network, once compromised, more frequently experiences compensatory mutation? Fig. 2 answers this question. As can be seen, the patterns of frequency of compensatory mutation depend on network size. For the smaller networks *N* = 5, 10, 15 and 20, the compensatory mutation rates continuously increase as network connectivity increases, but very gradually. In contrast, for the larger networks *N* = 30 and 40, with the rise in connectivity, the compensatory mutation rates decrease slightly. However, overall the results indicate that the frequency of individuals that can be fixed by compensatory mutation is more sensitive to network size than to network connectivity, and not particularly sensitive to either. The implied probability of compensatory mutation from the relative frequencies observed ranges from 8% to 15% of compromised networks recovering, with the larger rates associated with larger networks. This marked scale-invariance (see Fig. S1C, which is identical to Fig. 2 but re-scaled) stands in contrast to the scale dependencies shown for deleterious mutations in Fig. S1A and B.

**Figure 2:**
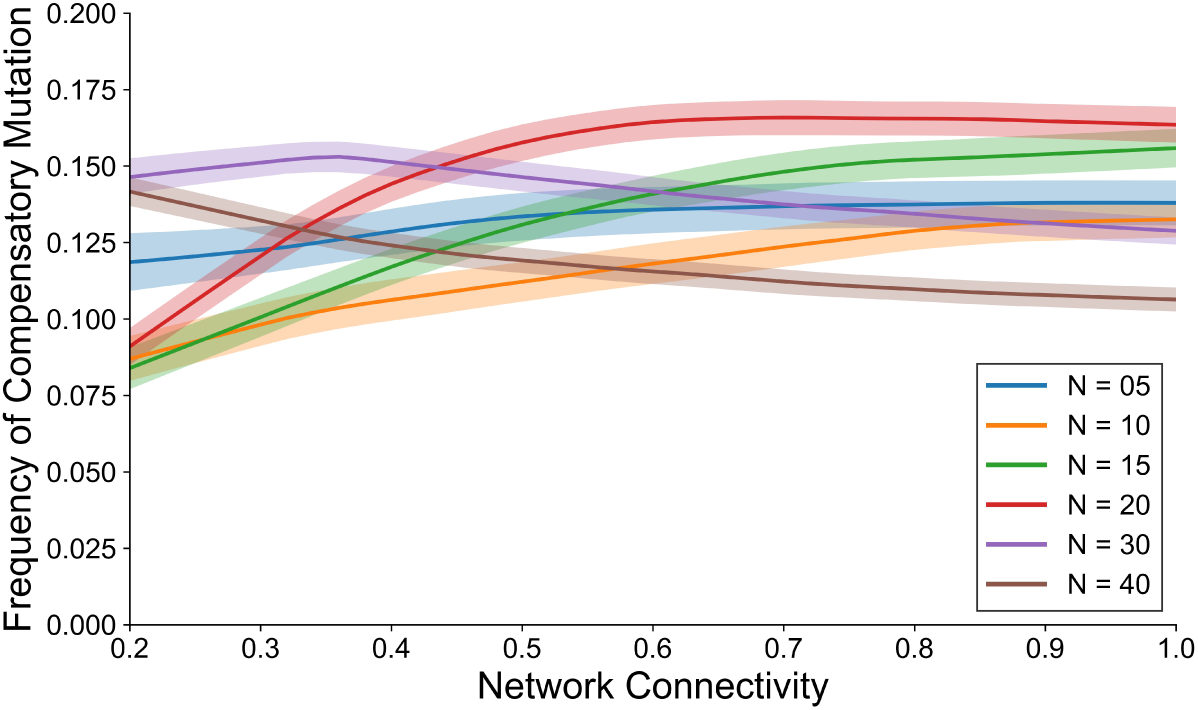
The frequency of compensatory mutation is relatively insensitive to network size and network connectivity in the context of gene regulatory networks. For each network size (*N* = 5, 10, 15, 20, 30 and 40 genes) for each value of network connectivity (proportion of regulatory relationships) given from a range of values in continuous intervals ([0.2, 1], step size 0.02), the frequency of experiencing first deleterious followed by a compensatory mutation on just two rounds of mutation was tested based on an initial 10, 000 randomly generated stable gene networks. The shaded areas represent 95% confidence intervals based on 100 independent runs.

Next, we investigate the occurrence of compensatory mutations in populations that have been exposed to bouts of generations of relaxed selection and selection for network stability. Again as a slight modification of what is standard for this type of model, we assume selection favours network stability. We find that compensatory mutation occurs in both evolutionary scenarios. From Fig. S2A, we can see that compensatory mutation is able to occur even in highly stable networks that have been subjected to network stability selection for many generations. In addition, the compensation probability tends to be constant after many rounds of mutation. Furthermore, we find that, across network sizes, all populations still maintain a high diversity in the presence of selection for network stability, and for many generations (see Appendix). It is not surprising therefore to see that, as shown in Fig. S2B, compensatory mutation can occur in the mixed populations (stable and unstable networks) that result from a relaxed selection regime, although it is less pronounced there and declines significantly over rounds of selection. Interestingly, we found that compensatory mutation can still fix seriously damaged networks that have suffered many deleterious mutations over generations. Here we select for study only the broken networks after each mutation round, as shown in Fig. S2C. Compensatory mutations will restore, for example, about 14% of networks for *N* = 5 that are broken by one round of mutation, but the frequency of compensation quickly drops to mutation restoring the stability of only 5% broken networks that have had many deleterious mutations for up to 15 generations. This indicates that compensatory mutations can still be cure-alls even for even seriously damaged networks, or at least those long neglected by selection for network stability.

Finally, we investigate the impact of relaxed selection on compensatory mutations. We find that, as expected, we can observe more compensatory mutations in the presence of relaxed selection for network stability. Specifically, we performed simulations to measure the number of compensatory mutations in lineages for which relaxed selection occurs in different likelihoods. From Fig. S3, we see that the number of compensatory mutations markedly increases as the consequence of having more generations of relaxed selection. We can also see that smaller networks typically have more compensatory mutations compared with larger networks. This is because compromised networks with smaller sizes are more likely to experience compensation after lengthy evolution, though larger networks tend to have a higher frequency of compensatory mutation at early stages as indicated in Fig. 2.

### Compensatory mutations exhibit bias in location and size

Given a model capable of generating compensatory mutation, we next characterise their nature. In general, there is little difference between a compensatory mutation and a deleterious mutation — in fact, the exact same mutation could be deleterious in one network and compensatory in another. However, compared to a completely random baseline, we find evidence that compensatory mutations tend to be biased with respect to the location in the network and the magnitude of gene regulatory effects.

We first look at where compensatory mutations happened in compromised networks. We find that they are more likely to occur at or close to the site of the original, deleterious mutation. In a typical small network with size *N* = 5 genes (see Fig. S4A), we found a 95.8% chance that a mutation that occurs on the exact site of a deleterious mutation compensates for it. The frequency of compensatory mutation is also high on most of the edges close to the original mutation site. Mutations on edges far away from the deleterious mutation site are much less likely to experience compensation. The same basic pattern is also seen in a larger network with size *N* = 20 genes (see Fig. S4B), where the frequency of mutations being compensatory, if they occur on the original deleterious site, is 85%. The percentages beside each edge in these figures indicate the proportion of mutations that occur on that edge that are compensatory, out of the 1, 000 simulated second rounds of mutation we ran on each edge for each network after it had previously suffered a single deleterious mutation. In general, as these representative figures indicate, the compensatory effect could happen in many positions in a broken network, but it is more likely to be observed at sites that are close to a deleterious mutation’s site.

Fig. 3A (solid line) demonstrates the generality of the result indicated in Fig. S4. It illustrates the frequency among 10, 000 initially-stable gene networks of compensatory mutation against different spatial distances from the single deleterious mutation suffered by each network. As can be seen, compensatory mutations generally occur in edges between genes close to the deleterious mutation site. We restrict the analysis to these five categories because there is only a narrow range of distribution distances for randomly sampled mutations (see Appendix for more details). We further conducted similar experiments for networks with neutral mutations to investigate whether compensatory mutations have any special property in terms of location. We found that, compared with the results of compensatory mutations, neutral mutations are more evenly distributed. Specifically, instead of measuring the frequency of a second, compensatory mutation (that restores network stability for a compromised network with a single deleterious mutation), we measure the frequency of a second, neutral mutation with different distance effects that retains the stability for a network that has already had one neutral mutation. From Fig. 3A (dashed line) we can see that the distance effect has a much less profound role in networks with two consecutive neutral mutations than in networks with one deleterious mutation and one compensatory mutation. In fact, neutral mutations tend to be enriched if they are far apart in larger networks (see Appendix).

**Figure 3:**
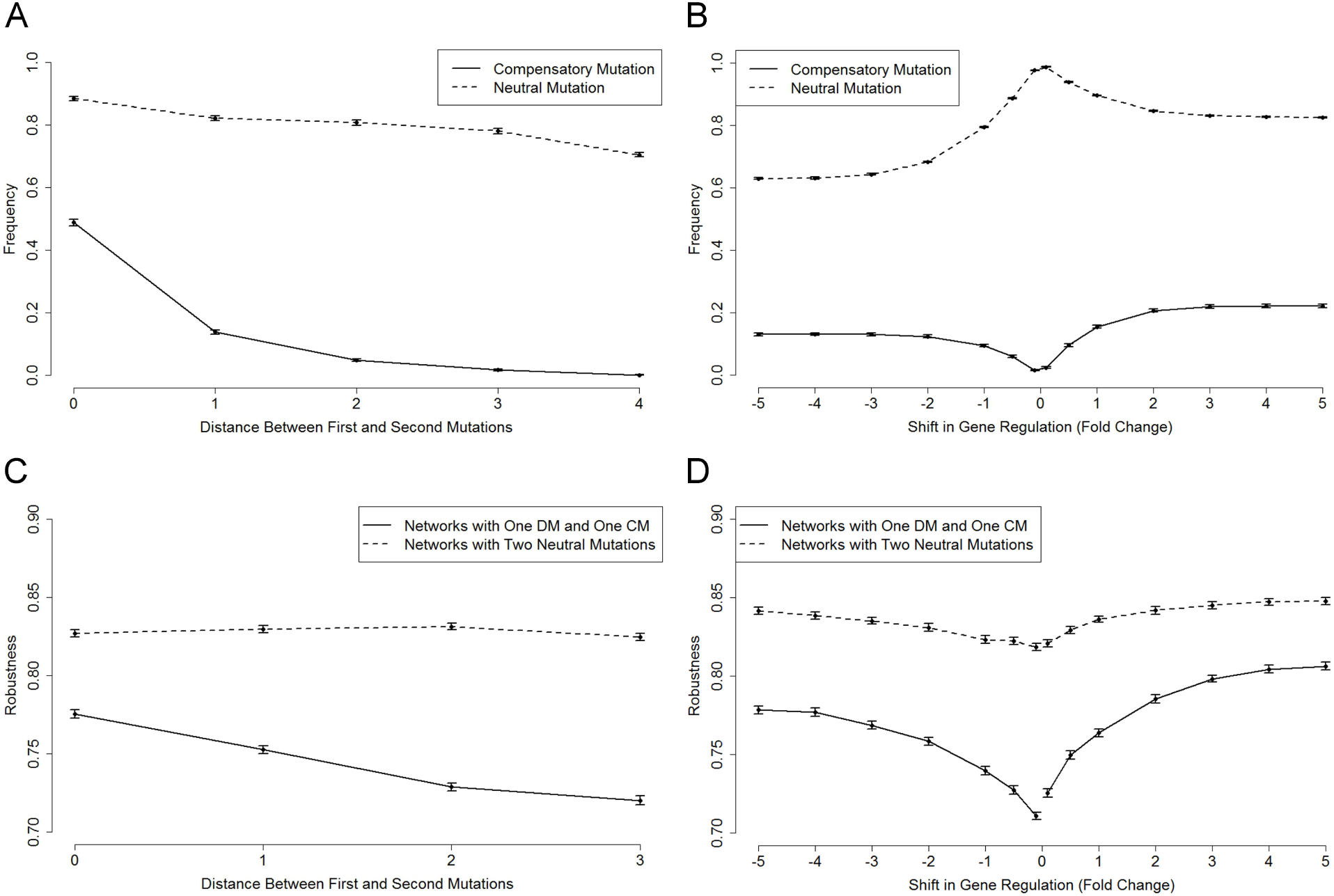
Both frequency and robustness of networks with compensatory mutations exhibit biases in location and mutation size. For *N* = 5 and *c* = 0.4, we first collected a pool of compromised networks with deleterious mutations after a single mutation round. We then forced second mutations, classifying these as being 0 (on the same site), 1, 2, 3 and 4 steps away from the original deleterious mutations (**A**) or adding a weight from [−5, +5] (step size 0.5) to the original regulatory impact (**B**). For each of these distance or weight categories, we measured the probability that the mutation was compensatory (that it returned the network to stability, see solid lines in (**A**) and (**B**)), based on 10, 000 sample networks collected for each category. The sample networks for control groups (see dashed lines in (**A**) and (**B**)) were collected in a similar way, except that the networks were subjected to two consecutive neutral mutations. Similarly, for *N* = 5 and *c* = 0.4, we collected 10, 000 sample stable networks that were subjected one deleterious mutation and then restored by one subsequent compensatory mutation that was 0, 1, 2 and 3 steps away from the previous deleterious mutation (**C**) or with different shifts in gene regulation from [−5, +5] (step size 1 and with four additional regulation shifts: 0.5, 0.1, 0.1 and 0.5) (**D**). Then, we assessed the robustness of the sample networks at each category (see solid lines in (**C**) and (**D**)). The sample networks for control groups (see dashed lines in (**C**) and (**D**)) were collected in a similar way, except that the networks were subjected to two consecutive neutral mutations. Error bars represent 95% confidence intervals based on 100 independent runs.

The point of compensation is of course to recover the network’s fitness, or here, stability. We therefore next investigate whether there is an impact on the robustness of networks after deleterious followed by compensatory mutation varies by the location of the compensatory mutation, and contrast this with differences in robustness after two neutral mutations. We again find that patterns are very dependent on location. Specifically, we compare robustness of stable networks following one round of deleterious and compensatory mutation with that of stable networks with two consecutive neutral mutations, as shown in Fig. 3C. In general, robustness is far higher when compensatory mutation occurs closer to the original deleterious mutation site (see the solid line in Fig. 3C), whereas after two neutral mutations, closer distances are not better associated with higher robustness (see the dashed line in Fig. 3C)^1^. Even though networks with compensatory mutations occurring near to the site of the deleterious mutation exhibit profoundly more robustness than those at other locations, their actual robustness is much lower than that of networks with neutral mutations (see Fig. 3C and Appendix). Nevertheless, these theoretical results indicate that these co-localised compensatory mutations are more likely to be accumulated, whereas compensatory mutations that are far apart from the previous deleterious mutations are more likely to be lost during subsequent selection for network stability.

Independent of location, we also investigated how different mutation size influences the probability of compensation in compromised networks. We found that compensation is more likely to be driven by large-effect mutations. Fig. 3B (solid line) presents the frequency of compensatory mutation against various intensities of up or down regulation among 10, 000 randomly-generated stable gene networks that had experienced a single deleterious mutation. For a randomly-chosen site in each network, we experimented with mutations across a range of regulatory strengths. As can be seen, larger regulation changes, both positive and negative, are up to a point associated with an increased frequency of compensatory mutation. However, the shape of the curve for compensatory mutations across all edges is not a symmetrical ‘V’. Rather, compensatory mutations occur more by positive changes to gene regulation than by negative changes. The explanation for this phenomenon is rooted in the fact that there are two edge types that can be affected by compensatory mutation: inter-gene regulation connecting two different genes and self-regulating edges. In the simulations, almost no compensatory mutations are both negative and self-regulating (see Appendix). The ‘V’ shape for only inter-gene regulation is almost symmetrical (see Appendix), suggesting that for these, negative and positive regulations are equally likely to be useful. It is true for both the negative and positive cases that compensatory mutation is increasingly likely with greater regulatory strength up to a certain extent.

Although we found that compensatory mutation tends to positive, this is not a property special only to compensatory mutations. From Fig. S5A, we can see that there is more positive regulation in both initially-stable networks and networks with compensatory mutations, whereas deleterious mutations in compromised networks tend to be more negative. By separating self- and non-self-regulatory edges, we find that compensatory mutations have a larger effect (in terms of shifting gene regulation) on self-regulatory edges than non-self-regulatory edges (see Fig. S5B and C). We then conduct similar experiments for networks with neutral mutations to investigate whether compensatory mutations have any special property in terms of mutation size. We find that, compared with the results of compensatory mutations, small-size mutations are more likely to be observed in networks with neutral mutations. Specifically, similar to the location experiments, we measured the frequency of a second mutation (neutral mutation) with different mutation effects that can retain the stability for a network that has already had one neutral mutation. From Fig. 3B (dashed line) we can see that that neutral mutations are more likely to be of small magnitude than large, and where they are large they are more likely to be positive than negative. Compensatory mutations are more likely to be of large magnitude than small, but are also more likely to be positive than negative.

As with location, we also investigated the robustness of networks subject to mutations of different sizes. We found that patterns of shifting regulation-generating robustness are also quite different. Specifically, we compared robustness of stable networks having one deleterious mutation and compensatory mutation with that of stable networks having two consecutive neutral mutations, as shown in Fig. 3D. In general, the robustness is significantly higher when compensatory mutation has a larger shift in gene regulation (see the solid line in Fig. 3D). Although networks with neutral mutations tend to have a similar pattern (see the dashed line in Fig. 3D), by measuring the percentage change in robustness (see Appendix), we can clearly see that size has a greater impact on robustness in compensatory mutations. Here again, it should also be noted that although networks with compensatory mutations exhibit a more profound biased change in robustness with respect to mutation size, their actual robustness is lower than that of networks with neutral mutations (see Fig. 3D). These theoretical results indicate that these large-effect compensatory mutations are more likely to be accumulated, whereas small-effect compensatory mutations are more likely to be lost by subsequent selection for network stability.

Note that similar patterns to those described here are also observed in networks with different sizes and connectivity. See more supporting figures in the Appendix.

### Compensatory mutation generates regulatory complexity

We now explore the long-term evolutionary consequences of compensatory mutations. Given the biases identified in the previous section concerning location and magnitude, we might predict that the effects of these two fundamental network properties would facilitate an altered neutral evolution, at least during periods of relaxed selection. Recall that we assume such relaxed periods will be interspersed between bouts of selection for network stability, and also that selection in our model favours network stability. We in fact do observe in our simulations an increase in the complexity of gene regulatory networks, but only in a context where they have been withdrawn from the selection for network stability for at least some proportion of generations. Specifically, we first generate a pool of 10, 000 stable networks (*N* = 10) with a simple ‘Star’ topology (see Fig. 4A), then evolve the population under different evolutionary scenarios. Fig. 4B shows four evolutionary scenarios where the population is exposed to selection for network stability in every generation such that there is no opportunity for compensatory mutation. From the typical results (networks with a median connectivity), we find that:

**Figure 4:**
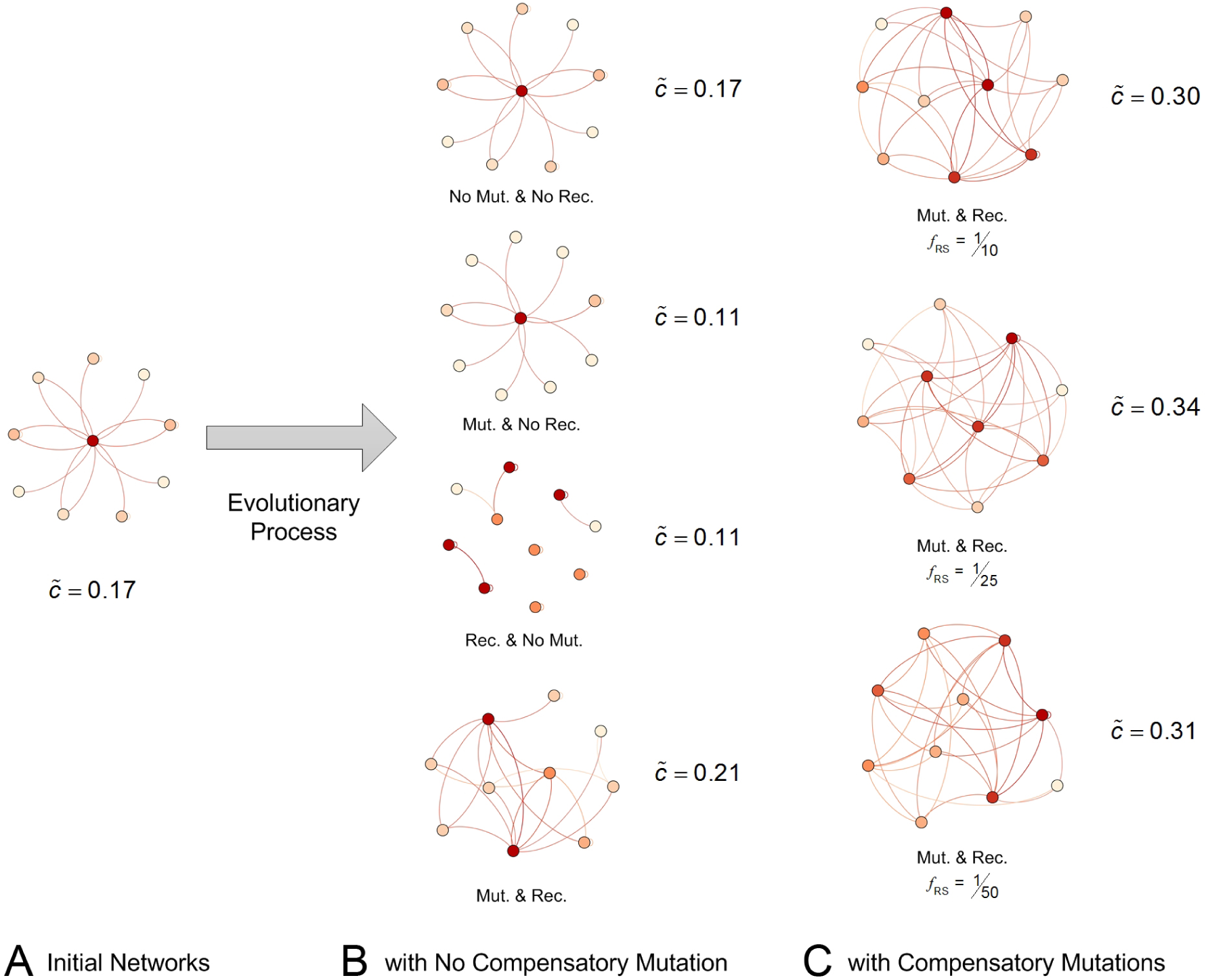
Compensatory mutation generates regulatory complexity in stable networks without an initial variation in network structure. The initial population pool was composed of 10, 000 sample stable networks with *N* = 10 genes. These networks had a similar “Star” topology (one hub node and nine non-hub nodes), varying network connectivity in [0.10, 0.26]. The detailed description of generating initial population can be found in Appendix. A representative network from the initial population is shown in (**A**). The initial population was evolved for 5000 generations with the selection for network stability (*f RS*= 0) under no mutation and no recombination regime, mutation but no recombination regime, recombination but no mutation regime, mutation and recombination regime (**B**). The initial population was also evolved for 5000 generations under relaxed selection regime (unstable networks will not be eliminated in generations under relaxed selection) with frequency *f_RS_* = 1*/*10, 1*/*25, 1*/*50 (**C**). Note that compensatory mutation cannot happen when the population is subject to selection for network stability, since no deleterious mutations survive to be compensated. The representative networks were selected randomly with a median connectivity in the evolved populations.

1. the median connectivity is the same as the initial population’s if it is evolved without mutation or recombination (only by drift),
2. the median connectivity decreases if evolved under either a mutation but no recombination regime or a recombination but no mutation regime (although the network structures are greatly altered when invoking only recombination), and
3. the median connectivity increases to an intermediate level if evolved under a regime allowing both mutation and recombination.

Fig. 4C shows three evolutionary scenarios where the population is evolved with periods of relaxed selection, invoking mutation (including compensatory mutation) and recombination. From these typical and individual results (networks with a median connectivity), we can see that the median connectivity greatly increases and is higher than in the case when the population is subjected exclusively to the selection for network stability so that no compensatory mutation can occur.

We would like to quantify the impact of relaxed selection on regulatory complexity. In another experiment, we collect 10, 000 stable networks and then evolve them for 5, 000 generations, allowing both mutation and recombination. From Fig. S6, we can see that if there is no relaxed selection at all, the mean connectivity of the population is highly preserved during evolution, whereas the network connectivity increases if we allow compensatory mutations to occur in periods of relaxed selection. It should be noted that in the first experiment, as shown in Fig. 4, we fix the network structure but vary the network connectivity in the initial population, whereas we fix the network connectivity but vary the network structure in this second experiment. These results demonstrate that selection for network stability where it impedes deleterious and compensatory mutations constricts complexity, whereas compensatory mutations contribute to regulatory complexity as a part of neutral process.

## 2 Discussion

Compensatory mutations have long been considered the primary means by which low-fitness lineages might be able to be restored to high fitness [Levin et al., 2000, Crawford et al., 2007, Meer et al., 2010]. More recently, Dunai et al. [2019] suggest compensatory mutation may account for robust adoption of costly traits that are of critical importance to an organism in certain circumstances, such as antibiotic resistance. Given that most mutations are believed to be deleterious at least initially, some such process would be essential for mutation to contribute to genetic innovation and evolution more generally. However, the extent of the role of compensatory mutations has often been considered to be negligible because they were considered to be highly improbable and therefore rare. As such, they have not been studied extensively, and many of their general properties have been unknown.

If the results presented here in simulation hold for *in vivo* regulatory networks, then compensation may be far more probable and frequent than had previously been anticipated. Our results indicate that gene networks may by their nature be surprisingly robust, such that a wide variety of alterations to a compromised network may effect its recovery. Significantly, we find that the frequency of compensatory mutation — unlike deleterious mutations — is relatively invariant to the size of the network. This may mean that iterations of deleterious and compensatory mutation play a far larger role in evolution than previously thought. Our results provide a new account for why compensation can be so rapid, and also show a significant impact of effect size, both of which have been observed in the laboratory [Moura de Sousa et al., 2017, Dunai et al., 2019]. In fact, there is some indication that quasi-deleterious mutations such as initially-costly mutations that promote antibiotic resistance, may produce a context wherein ‘compensatory’ mutations may find new adaptive fitness peaks, doing more therefore than mere compensation [Moura de Sousa et al., 2017].

Taylor et al. [2015] have shown that a regulatory network can be rapidly rewired through *de novo* compensatory mutation. In our simulations, we also observed that the compensatory mutation can facilitate regulatory complexity in terms of increasing network complexity (Fig. 4 and Fig. S6). Key in our simulations were periods where networks evolved under neutral processes driven by biases in compensatory mutation, since the regulatory complexity was not directly adaptively selected. Therefore, we believe compensatory mutations may be expected to play an essential role in driving regulatory complexity through neutral or non-adaptive processes.

Bouts of deleterious and compensatory mutations might well facilitate the transition of the regulatory network to new fitness peaks [Weinreich and Chao, 2005]. Compensatory mutations have been observed empirically to have a positive correlation with drug resistance mutations, where low-fitness lineages can create intrinsic selection pressure to mitigate their deleterious effects through compensatory mutations [Comas et al., 2012, Brandis et al., 2012, de Vos et al., 2013, Brandis and Hughes, 2013, Song et al., 2014, Dunai et al., 2019]. There is other supporting evidence that compensatory mutation can help the transition of lineages towards new fitness peaks [Martinez et al., 2014, Ivankov et al., 2014, Szamecz et al., 2014, Filteau et al., 2015]. Moreover, some studies also show that compensatory mutations can help increase plasmid stability, and thus facilitate adaptation [San Millan et al., 2014, Porter et al., 2015, Harrison et al., 2015]. Yet despite suggestions in the literature that peak shifts must occur through low-fitness genotypes [Wagner and Wright, 2007, Romero and Arnold, 2009, Olson-Manning et al., 2012, Osada and Akashi, 2012, Barreto and Burton, 2013], few theoretical studies have focused on how the formation of regulatory networks could be influenced by this process. We hope with our paper we have begun to redress this.

Historically, interactions in mutations *in vivo* have been considered hard to measure and the results usually have weak statistical significance [West et al., 1998, 1999], though see Moura de Sousa et al. [2017]. Where measurement is difficult, exploration of theoretical possibilities through simulation offers an ideal means to identify and test for logically-coherent scientific hypotheses and to discover unanticipated consequences of these. These unanticipated consequences are predictions arising logically from the hypotheses the model expresses — predictions that can then inform our search for evidence *in vivo* [Bryson et al., 2007]. The ability to observe and manipulate thousands of modelled individuals in a matter of hours allows for a systematic exploration of largely unknown theoretical territory. In this paper, the extension of the previous simulation approaches, while primarily conceptual, is potentially of great theoretical importance. Unlike the previous research seminal to our own [Wagner, 1996, Siegal and Bergman, 2002, Azevedo et al., 2006], we have been able to assess the probability and impact of compensatory mutations, providing important theoretical underpinnings to explain known laboratory outcomes.

The use of binary fitness outcomes (0/1 for unstable/stable networks) that are only periodically tested by selection for network stability (selection for network stability) is operationally quite useful. This allows us to avoid making unrealistic assumptions about the selection coefficient distribution, and to proceed on the assumption that moderately deleterious mutations may persist long enough to allow the accumulation of subsequent mutations, some of which may prove to be compensatory or even advantageous. Periodic assessment of the functional operation of networks, i.e., periods of selection for network stability, is still a necessary practical consideration. The fluctuating selection regime (periods of selection for network stability) modelled in this paper is certainly biologically realistic. For example, Siepielski et al. [2009b] conclude that selection is usually fluctuating following their study of the temporal dynamics of selection in a database which contains 5, 519 estimates of selection of wild populations. Similar arguments using empirical evidence can be found in Brachi et al. [2013], Gompert et al. [2014], Seppälä [2015] and Bijleveld et al. [2015].

Previous work has been taken to indicate that compensatory mutation is not likely to play an important role in the evolution of independently acting genes. The frequency of deleterious mutation is low; the frequency at which a new mutation compensates for the previous deleterious mutation had been expected to be even lower. However, mutations do not just happen in independently-acting genes. There is substantial molecular evidence for mutations in genes which exhibit complex interactions with other genes [Wilke and Adami, 2001, Wilke et al., 2003, Beerenwinkel et al., 2007, Lehner, 2011, Rokyta et al., 2011, Park and Lehner, 2013, Connelly et al., 2014]. In fact, gene regulatory networks are more likely to be able to accommodate deleterious mutations and therefore be available for compensatory mutations. The mutations simulated in models such as those presented here refer to mutations that occur in the binding sites of proteins at an enhancer, but not mutations in protein coding sequences. As such, their regulatory effects could be buffered by epigenetic neutrality, and evolve phenotypically neutral [Wagner, 1996, Espinosa-Soto et al., 2011]. The plasticity that evolves from such a system consequently increases the opportunity for compensatory mutations. The frequency at which deleterious mutations compromise gene regulatory pathways is likely to be substantially higher than that for an independently acting gene because there will inevitably be more possible sites to mutate. In this paper, we have demonstrated support for this possibility, that compensatory mutation could potentially be frequent (Fig. 2) and occur to some extent regardless of patterns of selection that the networks have been through (Fig. S2A and B). We have also shown that compensatory mutation can still occur even among seriously damaged networks (Fig. S2C). This is consistent with the findings of empirical studies, such as that by Sloan et al. [2014], who found that two *Silene* species with fast-evolving plastid and mitochondrial DNA exhibited increased amino acid sequence divergence in organelle genomes but not in cytosolic ribosomes. Given that the authors found no evidence that the observed pattern was driven by positive selection, they concluded that the rapid organelle genome evolution had selected for compensatory mutations in nuclear-encoded proteins. More recently, both Moura de Sousa et al. [2017] and Dunai et al. [2019] find that compensatory mutation rather than reversion are actually fairly outcomes for bacteria developing resistance to multiple antibiotics at once. In our present paper, we have demonstrated theoretical support and explanation for these empirical findings, by showing that compensatory mutations can be greatly increased if the population evolves during a phase of relaxed selection regime (Fig. S2). Here, ‘relaxed’ is more obviously a relative term. Organisms challenged by antibiotics are already stressed, and trade-off the costs of initial mutations with the benefits in surviving the antibiotic assault. In this climate, ‘compensatory’ mutations are mutations selected because they mitigate these additional costs, reducing the chance that the mutations leading to antibiotic resistance are swept through reversion from the population. Although antibiotic resistance is obviously a problem in human contexts, more generally this sort of dynamic illustrates a context in which ‘relaxed’ selection might come into play — when a mutation produces both costs *and* benefits, or these vary with the ecological context of the organism.

Many studies have shown that conventional *de novo* mutations are widely distributed throughout the genome and have a wide distribution of phenotypic effects, from complete lethality to weak benefit with respect to fitness [Sanjuán et al., 2004, Eyre-Walker and Keightley, 2007, Keightley and Eyre-Walker, 2007, Mezmouk and Ross-Ibarra, 2014]. Although there have been no predictive tests of the location of compensatory mutations, empirical studies show that compensatory mutations are often found in proteins that are in or interact with proteins that exhibit a deleterious mutation [Poon et al., 2005, Poon and Chao, 2005, Davis et al., 2009, Comas et al., 2012, Bhattacherjee et al., 2015]. Our findings concur with this. In this paper, we have showed that there is a bias with respect to where compensatory mutations happen such that compensatory mutations tend to generate regulatory circuits that closely interact with each other (solid line, Fig. 3A), whereas neutral mutations tend to accumulate more evenly distributed and therefore further apart from each other (dashed line, Fig. 3A). We also found a bias with respect to the size compensatory mutations have in terms of shifting gene regulation, such that compensatory mutations generate regulatory circuits that have larger interactive impacts (solid line, Fig. 3B), compared to neutral mutations (dashed line, Fig. 3B).

Previous work has indicated that the origin of mutational robustness may come from the non-adaptive results of biophysical principles or non-adaptive evolutionary forces [Ruths and Nakhleh, 2013, Payne and Wagner, 2015]. During periods of relaxed selection, regulatory networks with otherwise-lethal mutations have the potential to be compensated by additional mutations. If compensatory mutation occurs frequently enough and generates different patterns of gene regulation than networks with neutral mutations, then the processes observed here could alter which types of network are lost when selection for network stability does occur. Systematic biases in the loss of particular network configurations could allow network features associated with compensatory mutation to accumulate in the population, even when the features do not at least initially confer differential reproductive success. In addition—as we have shown—the combination of recombination, deleterious mutation and compensatory mutation under moderately effective population sizes could then permit the evolution of increased regulatory complexity. In this paper, we have shown that stable networks with compensatory mutations generating a profound change in robustness compared to the impact on stable networks of neutral mutations (Fig. 3C and D). These results indicate that over time, compensatory mutations that occur during generations of relaxed selection could be biased such that regulatory circuits that closely interact and have larger interactive impacts are more likely to be maintained. We have also shown that, at least in our system, over time these can have profound impact on the complexity of the networks.

Taken together, we believe these findings demonstrate that the nature of compensatory mutation has been misunderstood theoretically. Periods of relaxed selection (as per our model) or indeed of increased differential selection (as per ecological challenges e.g. of antibiotics) produce a context in which natural innovations may be tolerated long enough to be combined. Combining mutations allows for a larger range of genomic innovation. Both our models and the empirical data of others show a surprising level of resilience in complex biological systems. Our models indicate that resilience may scales well with increasing complexity. Overall, the biases that emerge in this process as innovations accumulate may be an important new factor in understanding the evolution of gene regulatory networks, and evolution more broadly.

## Methods

We employed a well-established synthetic model of gene regulatory networks to simulate compensatory mutation; see Appendix for further in-depth description of our simulations. Here we only provide a more detailed explanation of the computational model used in this paper.

For each individual in a finite population of size *M*, we consider an *N × N* matrix *W* as a gene network that contains the regulatory interactions among *N* genes. Each element *w_i,j_* (*i, j* = 1, 2*, …, N*) represents the regulatory effect on the expression of gene *i* of the product of gene *j*. The network connectivity parameter *c* determines the proportion of non-zero elements in the network *W*. A zero entry means there is no interaction between two genes. Through gene interactions, the regulatory effect acts on each gene expression pattern. This can be denoted by a state vector **S**(*t*) = (*s*_1_(*t*)*, s*_2_(*t*)*, …, s_i_*(*t*)*, …, s_N_* (*t*)), where *s_i_*(*t*) represents the expression level of gene (or concentrations of proteins) *i* at time *t*. Each value of expression state *s_i_*(*t*) is within the interval [*−*1, +1] that expresses complete repression (*−*1) and complete activation (+1). For a given gene regulatory network *W*, the dynamics of **S** for each gene *i* is modelled by a set of coupled difference equations:

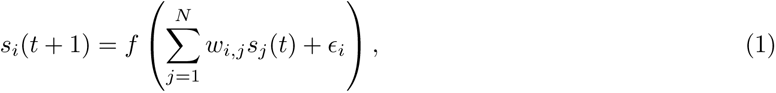

where *f* (*·*) is a sigmoidal function, and *E_i_* is a constant which reflects either a basal transcription rate of gene *i* or influences of upstream gene(s) on gene *i*. For reasons of c omputational c onvenience, we set *E _i_* = 0, and follow Siegal and Bergman [2002] and Azevedo et al. [2006] to define *f* (*x*) = 2/(1 + *e^−ax^*) *−* 1, where *a* is the activation constant determining the rate of change from complete repression to complete activation.

In all the simulations here, we define network developmental stability as the progression from an arbitrary initial expression state, **S**(0), to an equilibrium expression state (reaching a fixed phenotypic pattern), **S**_EQ_, by iterating Equation (1) a fixed number of times, *devT*. If a given network *W* can achieve stability over this developmental time period, it is termed ‘stable’; otherwise, it is labelled ‘unstable’. Note that this selection for network stability is also referred to as selection for network stability in which unstable networks will be eliminated. The equilibrium expression state can be reached when the following equation is met:

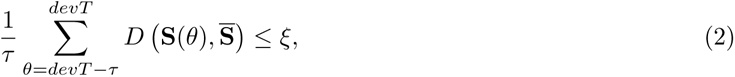

where *ξ* is a small positive integer and set to be 10*^−^*^4^ in all simulations, and 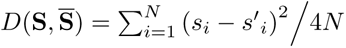 measures the difference between gene expression patterns **S** and 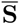 which is the average of the gene expression level over the time interval [*devT − τ, devT − τ* + 1*, …, devT*], where *τ* is a time-constant characteristic for the developmental process under consideration, and depends on biochemical parameters, such as the rate of transcription or the time necessary to export mRNA into the cytoplasm for translation [Wagner, 1994]. Unless otherwise specified, we used *a* = 100, *devT* = 100 and *τ* = 10 in all simulations, following previous studies [Wagner, 1996, Siegal and Bergman, 2002, Azevedo et al., 2006].

### Initialisation

Each individual network in the population was generated with a gene regulatory matrix *W* associated with an expression state vector **S**(0). Specifically, the matrix was generated by randomly filling *W* with *c × N* ^2^ non-zero elements *w_i,j_* that was drawn from a standard normal distribution *N* (0, 1). The associated initial expression state **S**(0) was also set by randomly choosing each *s_i_*(0) = +1 or *−*1.

### Mutation

In the mutation operation, exactly one element *w_i,j_* picked at random in each regulatory matrix *W* would be replaced by *w′_i,j_ ∼ N* (0, 1). Note that the mutation only occurs among non-zero elements. In other words, the mutation process will not change the topology of the original network *W* in terms of forming new edges or deleting existing edges between two genes.

### Recombination

In some simulations presented in this paper, we allowed individual networks to recombine with each other. A recombinant was produced by picking two individuals and selecting rows of the *W* matrices from each parent with an equal probability. This process is similar to free recombination between units formed by each gene and its *cis*-regulatory elements, but with no recombination within regulatory regions.

### Strong and relaxed selection for network stability

In the selection for network stability regime, only individuals which were able to attain developmental stability after the mutation process were selected. In contrast, all individuals can survive regardless of they were capable or incapable of reaching equilibrium when the selection for network stability was relaxed.

### Evolution

The evolutionary simulations were performed under the reproduction-mutation-selection life cycle. The population size *M* was fixed in every generation throughout the evolution in all simulations. In typical asexual evolution, an individual was chosen at random to reproduce asexually by cloning itself and was then subjected to a single mutation. Similarly, in typical sexual evolution, two individuals were chosen at random to reproduce sexually by recombining two parent networks and then subjected to a single mutation. Depending on different patterns of selection, unstable networks were excluded (under the selection for network stability regime) or allowed to stay in the population (under the relaxed selection regime). This process was repeated until *M* number of networks were produced.

## Data Availability

Simulation code for the simulations is available at https://bit.ly/2ExLhYd.

## Acknowledgements

Thanks to Alistair Fletcher for his early exploration of these models as part of his computer science masters degree at the University of Bath. The two final authors (JJB and NKP) are coequal.

## Supplementary Figures

**Figure S1:**
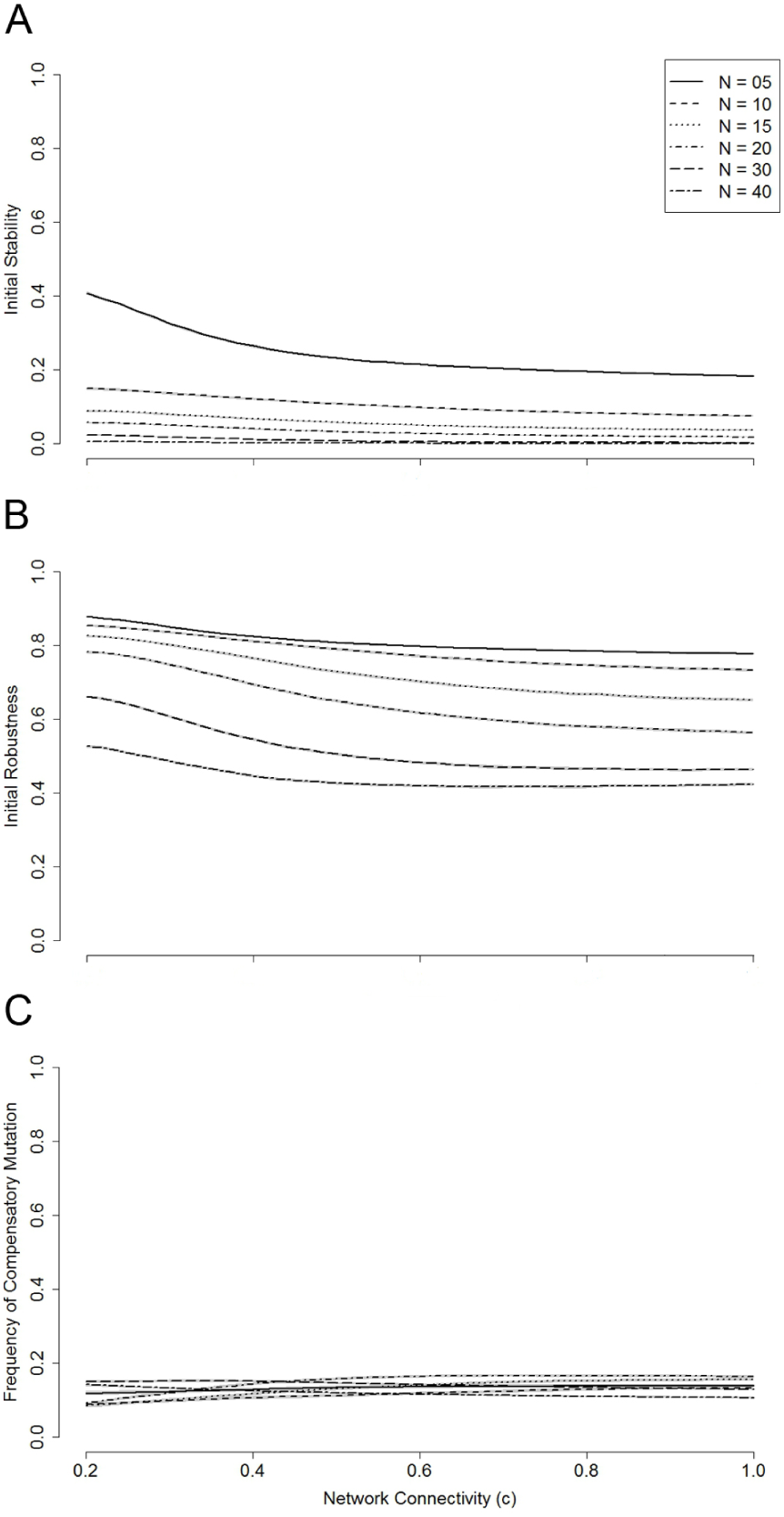
The influence of the size and connectivity of a gene regulatory network on its initial stability, robustness and frequency of compensatory mutation. For each network size (*N* = 5, 10, 15, 20, 30 and 40) with each connectivity given from a range of values in continuous intervals ([0.2, 1], step size 0.02), we tested the proportion of gene networks that are stable based on an initial 10, 000 randomly generated networks (**A**), the robustness of stable networks after exposure to a single round of mutation based on an initial 10, 000 randomly generated stable networks (**B**), and the frequency of compensatory mutation based on an initial 10, 000 randomly generated stable networks (**C**) (rescaled from Fig. 2). The shaded areas represent 95% confidence intervals based on 100 independent runs.

**Figure S2:**
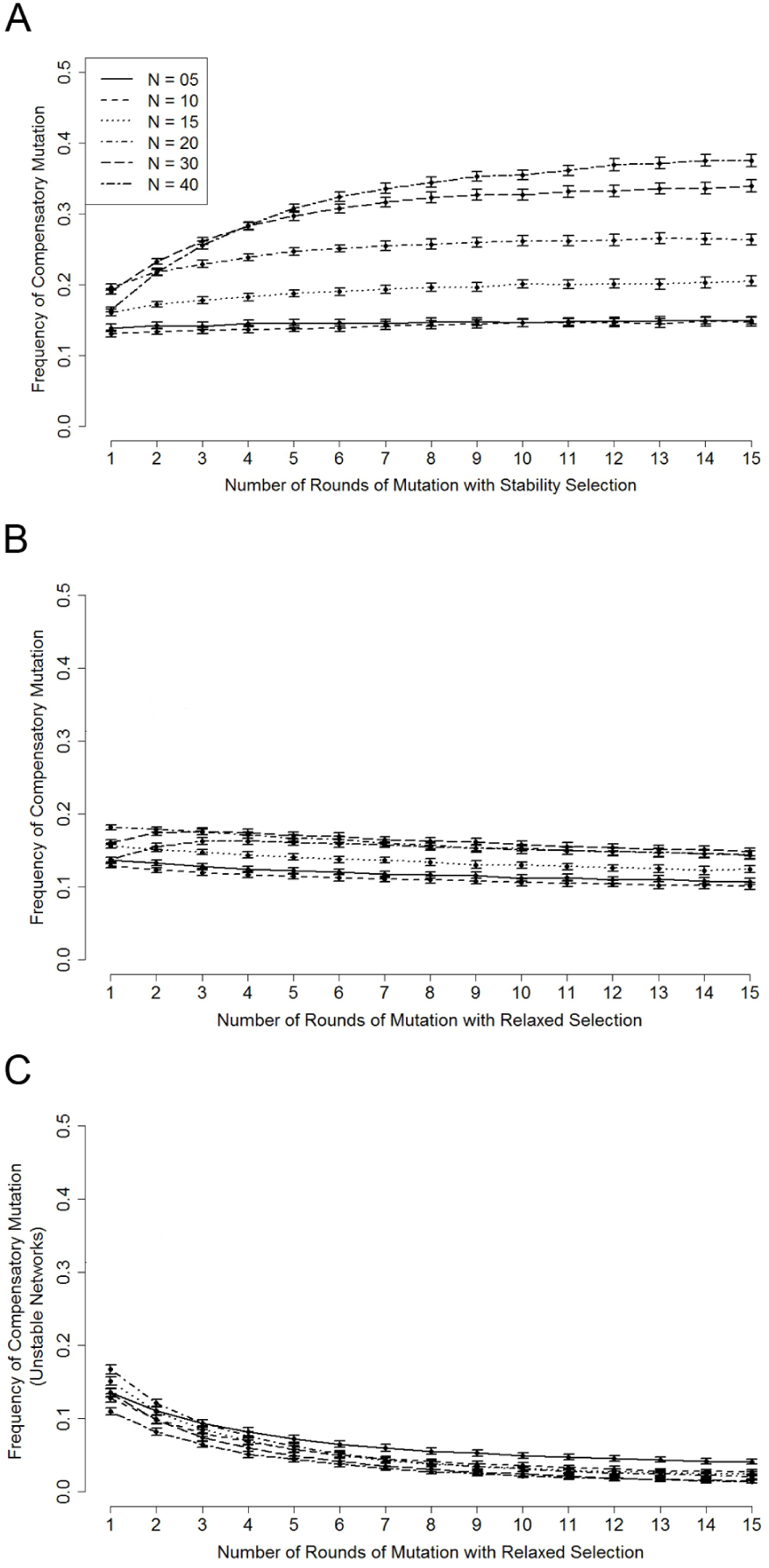
Compensatory mutations can occur in networks regardless of different patterns of selection. For each network size (*N* = 5, 10, 15, 20, 30 and 40) with network connectivity *c* = 0.76, we collected 10, 000 only stable, both stable and unstable, and only unstable networks with one to fifteen rounds of mutation. For each round of mutation, each network was subjected to one single mutation (for unstable networks) or two single mutations (for stable networks). Then, we measured the frequency of compensatory mutation in networks that have been subjected to bouts of selection for network stability (**A**), networks that have been subjected to bouts of relaxed selection (**B**), and networks with cumulative deleterious mutations (**C**). The error bars represent 95% confidence intervals based on 100 independent runs.

**Figure S3:**
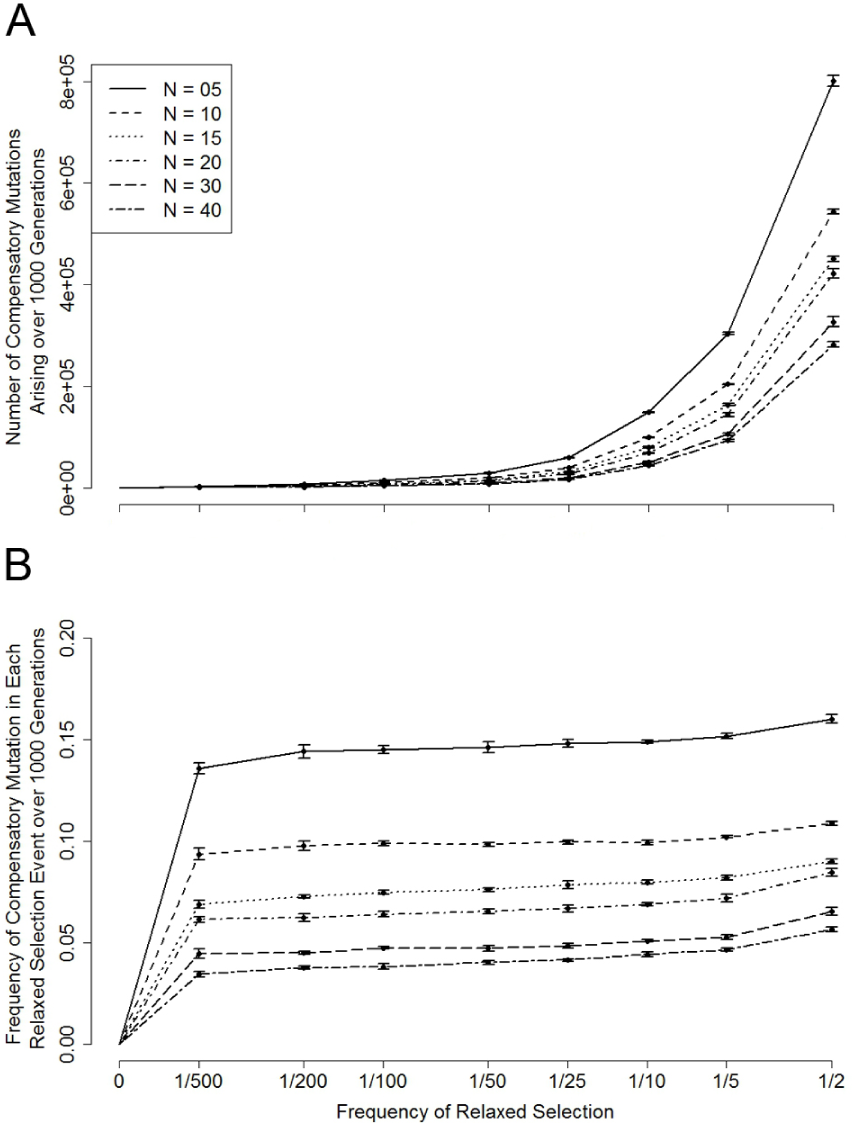
Relaxed selection stimulates compensation in gene regulatory networks. For each network size (*N* = 5, 15, 10, 20, 30 and 40) with connectivity *c* = 0.76, we measured the number of compensatory mutations occurring after the previous relaxed selection, which happened in every 2, 5, 10, 25, 50, 100, 200 and 500 generations. The reported results are the total number of compensatory mutations (of 10, 000 networks) (**A**) and frequency of compensatory mutation (per network per relaxed selection cycle) occurring over a total of 1, 000 generations for populations with different sizes (**B**). Error bars or shaded areas represent 95% confidence intervals based on 100 independent runs.

**Figure S4:**
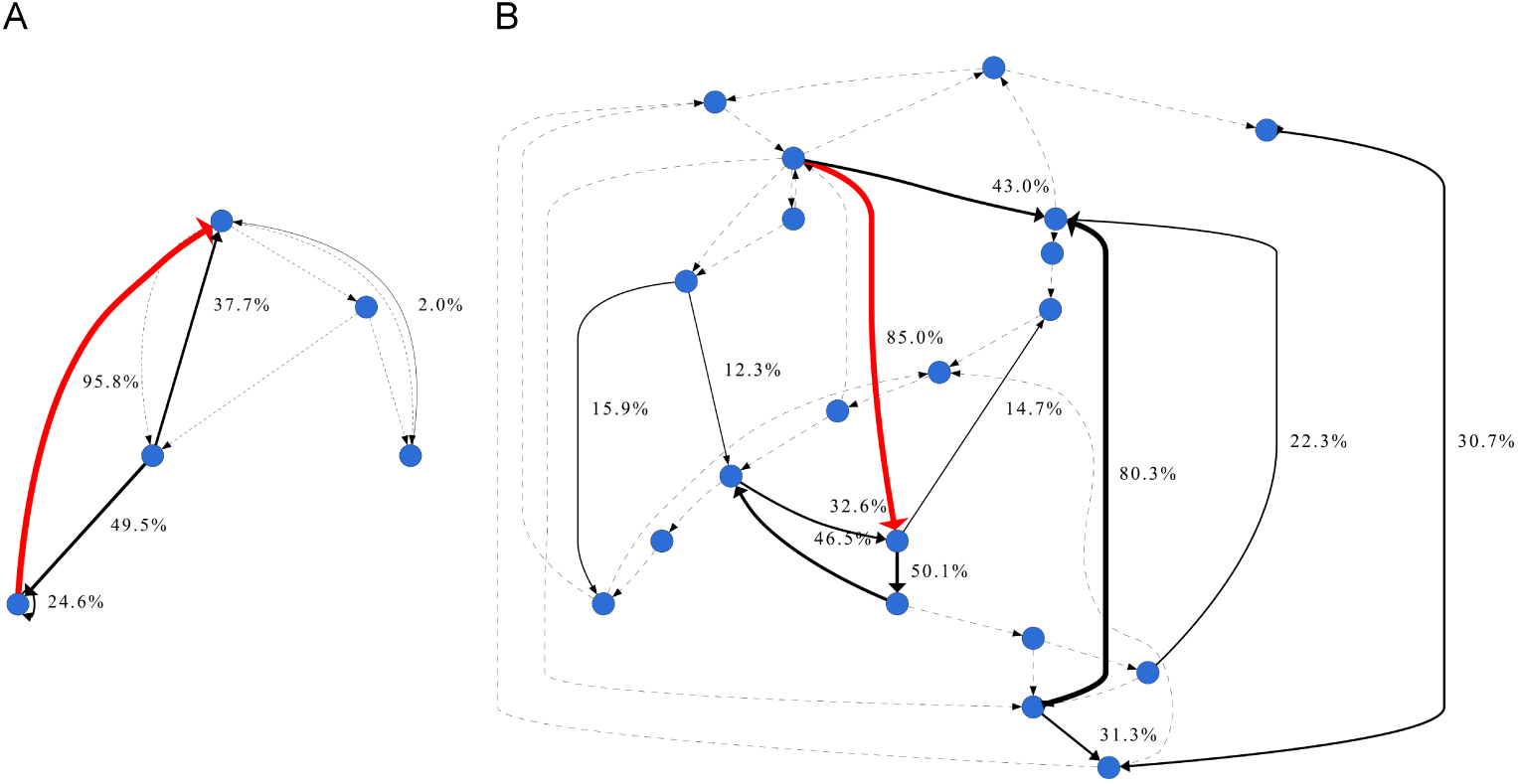
Examples of the spatial probability of compensatory mutation occurring on gene networks. In both examples, *N* = 5 (**A**) and 20 (**B**), for a particular compromised network that was stable initially, we executed one additional mutation round 1, 000 times on each edge. Then, the percentage of each mutation on that edge that restored GRN stability after mutation was measured. Unmarked edges had a CM 0% of the time. Note the solid line with width also indicates the probability an edge’s mutation was compensatory, and the dashed line to represent the edges for which mutation never compensated this particular deleterious mutation. The original deleterious mutation occurred on the edge marked in red. Note: The directed edge represents the interaction between two connected genes. But we do not distinguish negative or positive regulation in the provided examples.

**Figure S5:**
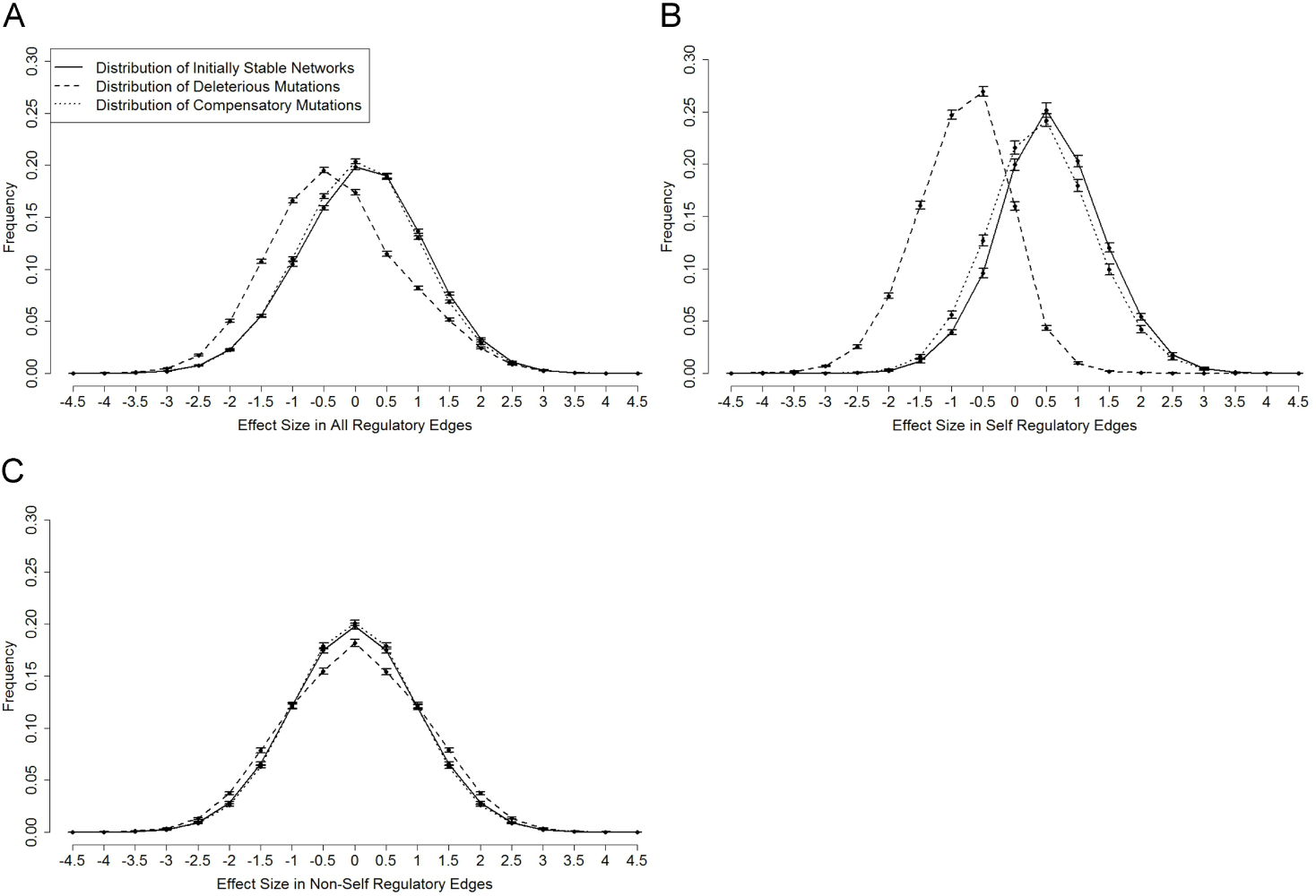
The distribution of regulation in initially-stable, compromised and restored networks. For randomly generated stable networks with *N* = 5 and *c* = 0.4, we collected 10, 000 sample regulations. We also collected 10, 000 sample regulation weights from deleterious mutations that compromised initially-stable networks as well as from compensatory mutations that restored the stability of previously-broken networks. We then measured the distributions in all regulatory edges (**A**), in self-regulatory edges (**B**) and ignoring self-regulatory edges (**C**). Given that the regulations are continuous values, we grouped them into 19 bins from [−4.5, +4.5] (step size 0.5). The error bars represent 95% confidence intervals based on 100 independent runs.

**Figure S6:**
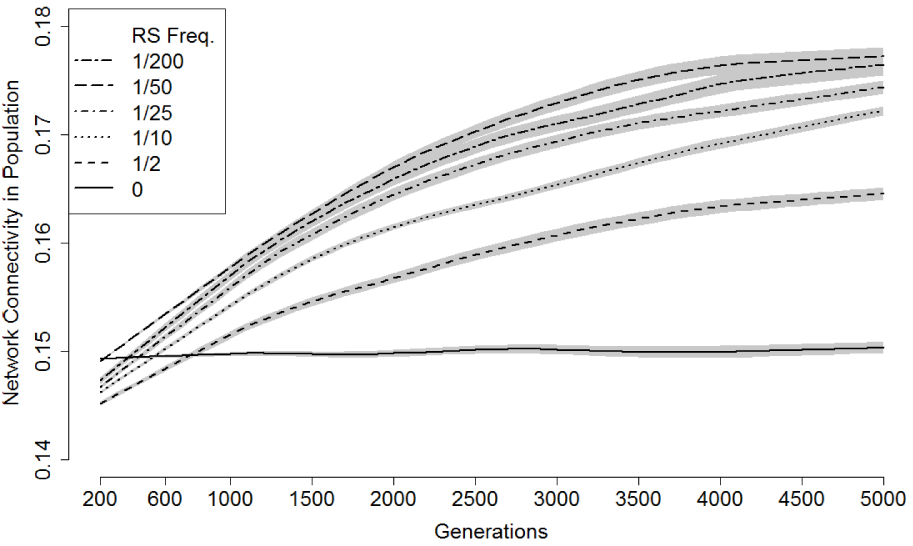
Compensatory mutation generates regulatory complexity in stable networks without an initial variation in connectivity. For the network size *N* = 40 and the connectivity *c* = 0.15, we collected 10, 000 stable networks, then evolved them for 5000 generations, allowing recombination at each generation. In every 200 generations, we measured the network connectivity of the population (stable) in which the relaxed selection occurs in every 2, 10, 25 50 and 200 generations. We also measured the network connectivity of the population when there was no relaxed selection as a control group. Shaded areas represent 95% confidence intervals based on 10 independent runs.

## Supplementary Text

### Estimating the relative frequency of compensatory mutation

To gain an impression of the properties of the initial gene regulatory networks, we first tested the probability of network stability in randomly-generated networks. As illustrated in Fig. S1A, smaller networks are more likely to be stable. Moreover, the relative frequency of stability in networks with low levels of connectivity is higher than that of networks with high levels of connectivity. This is in general accordance with previous work, typically done at connectivity *c* = 0.75, e.g. Azevedo et al. [2006], which indicates that larger networks with complex topology tend to be unstable.

In the second experiment, we explored the robustness of initially-stable networks; that is, we investigated the probability that stable networks remain stable after a single round of mutation. Here, a single mutation means exactly one non-zero entry in an individual’s genotype would be mutated. Given that the initially-stable networks were collected from the original randomly-generated ones, it would seem reasonable to predict that the small stable networks are more likely to break after one mutation round, since they contain fewer pathways and a single mutation, therefore, has a greater proportional effect. However, the results in Fig. S1B show the opposite effect: the stability of the small networks is still high. The mutation operation is effectively an alternative way of generating new networks; thus, the mutated networks have the same properties as the initial ones.

In our third experiment, we measured the compensatory mutation frequency in previously-stable networks. Specifically, we started from a population pool where each stable network was randomly generated. Then, we exposed these initially-stable networks to a single round of mutation. We focused on those unstable networks where each network contained a single deleterious mutation. Next, we exposed these compromised networks to an additional round of mutation. Finally, we tested the stability of the resulting networks. The stable networks at this point had experienced compensatory mutation. We then measured the frequency of individuals that experienced compensatory mutation. As shown in Fig. S1C (also see Fig. 2, main text), the frequency of compensatory mutation is largely scale invariant both to the network size and the network connectivity.

### Exploring strong and relaxed selection for network stability on compensatory mutation frequency

In this set of experiments, we investigated the frequency of compensatory mutation after many generations of both strong and relaxed selection for network stability to test whether compensatory mutation continues to occur even after lengthy evolution (see Fig. S2A and Fig. S2B). Specifically, under the selection for network stability regime, we collected 10, 000 stable networks at each generation where each network in the population was subjected to one single mutation. Then, we performed another round of mutation, focusing on the unstable networks that resulted from the previous round, and measured the probability of a second mutation that can restore the network stability of those compromised networks. Similarly, under the relaxed selection regime, we collected 10, 000 networks at each generation where each network in the population was subjected to one single mutation. However, for each relaxed selection generation, there were both stable and unstable networks after the population was subjected to the single mutation, since we did not perform selection restricting networks to being stable. The overall frequency of compensatory mutation for the population during each relaxed selection generation was averaged over the results of stable networks and unstable networks that were calculated separately.

In addition, we also measured the frequency of compensatory mutation among unstable networks during each relaxed selection event to further confirm that compensatory mutation can occur even in seriously damaged networks (see Fig. S2C). Specifically, we collected 10, 000 unstable networks at each generation where each network in the population was subjected to one single mutation, so really in this case we had selected against network stability. Then, we performed another round of mutations and measured the probability of a second mutation that could restore network stability. Note that this set of experiments is similar to those experiments described above, but here we only focus on unstable networks, whereas we consider both stable and unstable networks in the relaxed selection regime.

### Exploring the frequency of relaxed selection in simulating compensatory mutations

In this set of experiments, we tested whether frequent relaxed selection can generate more compensatory mutations (see Fig. S3A). Specifically, we collected a population pool of 10, 000 stable networks that were generated randomly. The initial population was then evolved under a relaxed selection regime with a frequency of 1*/*2, 1*/*5, 1*/*10, 1*/*25, 1*/*100, 1*/*200 and 1*/*500 for a total of 1, 000 generations. Note that during relaxed selection event, both stable and unstable networks can survive when the population is subjected to one single round of mutation. The number of compensatory mutations was recorded immediately after each relaxed selection event when the population was subjected to another single round of mutation. The reported results are the total number of compensatory mutations (see Fig. S3A) and frequency of compensatory mutation (per network per relaxed selection event, see Fig. S3B) arising over 1, 000 generations.

### Exploring population diversity for highly stable networks

In this set of experiments, we investigated how the population diversity is impacted in networks that have been exposed to many generations of strong selection for network stability (see Fig. S7). Specifically, we tested whether the increased compensatory mutation frequency shown in Fig. S2A was due to the property of particular networks that had been selected for, or whether it was the property of a diverse population. Following the measurement used in Azevedo et al. [2006], the genetic diversity is defined as:

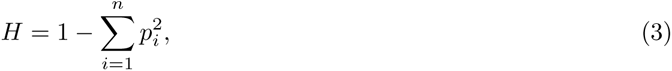

where *n* is the total number of alleles, i.e., the unique values contained in the same site crossing all individual networks, and *p_i_* is the frequency of allele *i*. The genetic variation in a population is calculated as the mean gene diversity over non-zero sites of the interaction matrix for a given genotype *W*.

**Figure S7:**
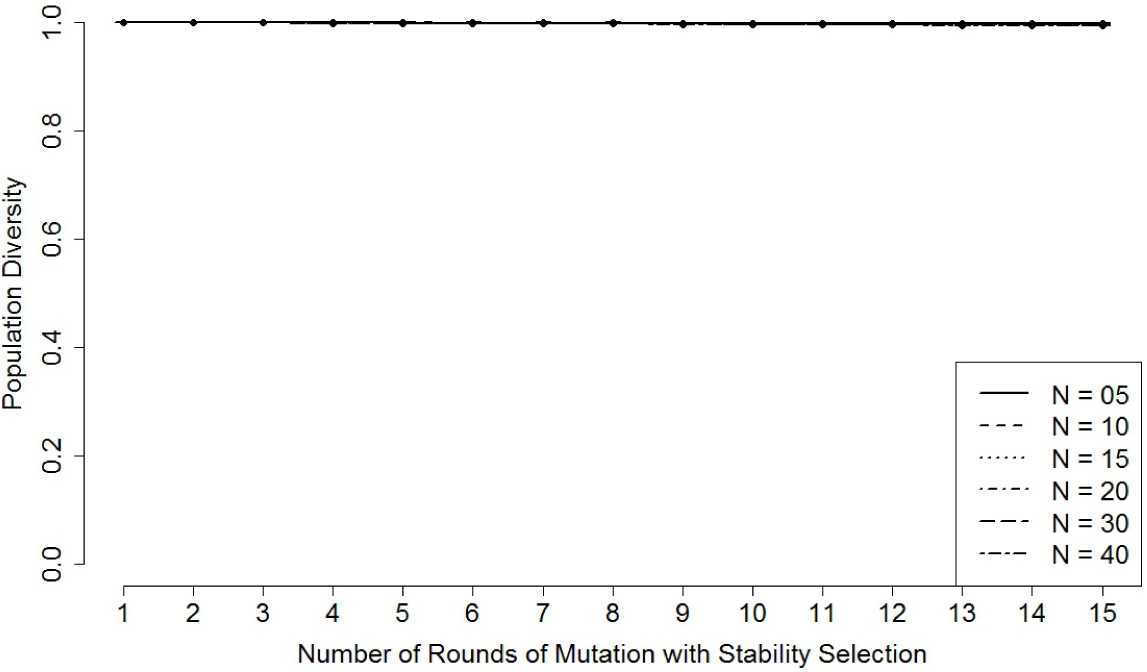
Population diversity of highly stable networks. For each network size (*N* = 5, 10, 15, 20, 30 and 40) with network connectivity *c* = 0.76, we tested population diversity for 10, 000 networks that had been exposed to selection for network stability following up to fifteen rounds of mutation as described in Fig. S2A. The error bars represent 95% confidence intervals based on 100 independent runs.

From Fig. S7, we can see that networks that have been though many generations of selection for network stability can still maintain a high network diversity.

### Location of compensatory mutations

In this set of experiments, we first sought to visualise locations at which the compensatory mutations are more likely to occur (see Fig. S4). To this end, in a set of compromised networks (those stable networks that proved fragile to a single round of mutation), we marked the site of the deleterious mutation, then measured the relative frequency of compensatory mutation that occurred at each possible site, including the site of the deleterious mutation, within this compromised network. For each possible site, we measured the outcomes over 1, 000 simulated mutations on that site (so that only the extent of regulation was mutated randomly, not the location).

To quantify the distance between deleterious and (potentially) compensatory mutation, we first define ‘distance’ as used in this paper. Suppose a given gene regulatory network, denoted as *W*, has two marked edges denoted as 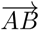 (deleterious mutation) and 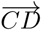 (compensatory mutation), where *A*, *B*, *C* and *D* represent different genes in *W* and 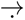 marks the edge direction. The distance between 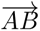 and 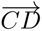 can be calculated as

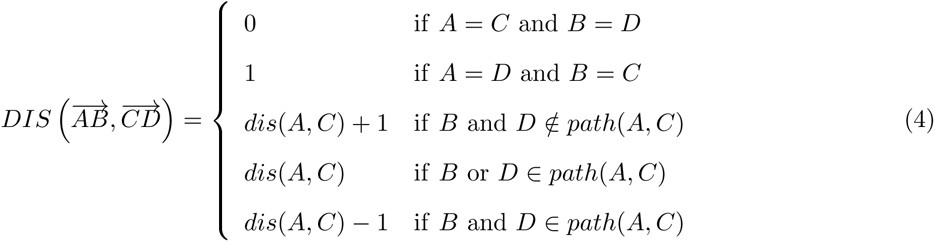

where *dis*(*A, C*) is the fewest edges possible from *A* to *C* and *path*(*A, C*) includes the vertices on the shortest path between *A* and *C* in network *W*.

An example process of compensatory mutation in a gene regulatory network can be seen in Fig. S8. This stable network can be compromised by a single deleterious mutation (marked in red) and compensated by an additional mutation (marked in blue). According to Equation (4), the distance from deleterious mutation site 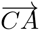 to compensatory mutation site 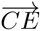 can be calculated as: 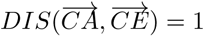

Next, we compared the relative frequencies of compensatory mutation among gene networks whose marked edges (caused by additional mutation) were 0, 1, 2, 3, and 4 steps away from the deleterious mutation (see Fig. 3, main text, and also see Fig. S9). We also performed similar experiments for medium (*N* = 20) and large networks (*N* = 40), as shown in Fig. S10 and Fig. S11.

**Figure S8:**
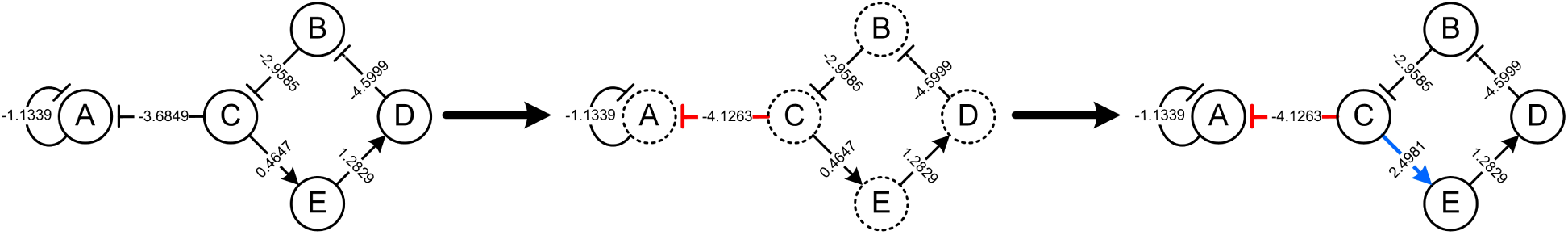
An example process of compensatory mutation in a gene regulatory network. The initially stable gene network contains five genes: *A*, *B*, *C*, *D* a nd *E*. I n t he i nitial n etwork (on t he l eft side), each directional edge represents the strength (weight) of interaction between the linked two genes. The initial gene expression pattern is **s**(0) = (1, 1, +1, +1, +1). In the compromised network (in the middle), a mutation occurs on 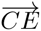 (indicated in red), which leads to the failure of stabilising the gene expression patterns (marked by dashed circles). In the compensated network (on the right side), the compromised network is fixed by an additional mutation that occurs on 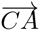 (indicated in blue), reaching an equilibrium expression **s**_EQ_ = (*−*1*, −*1, +1, +1, +1).

**Figure S9:**
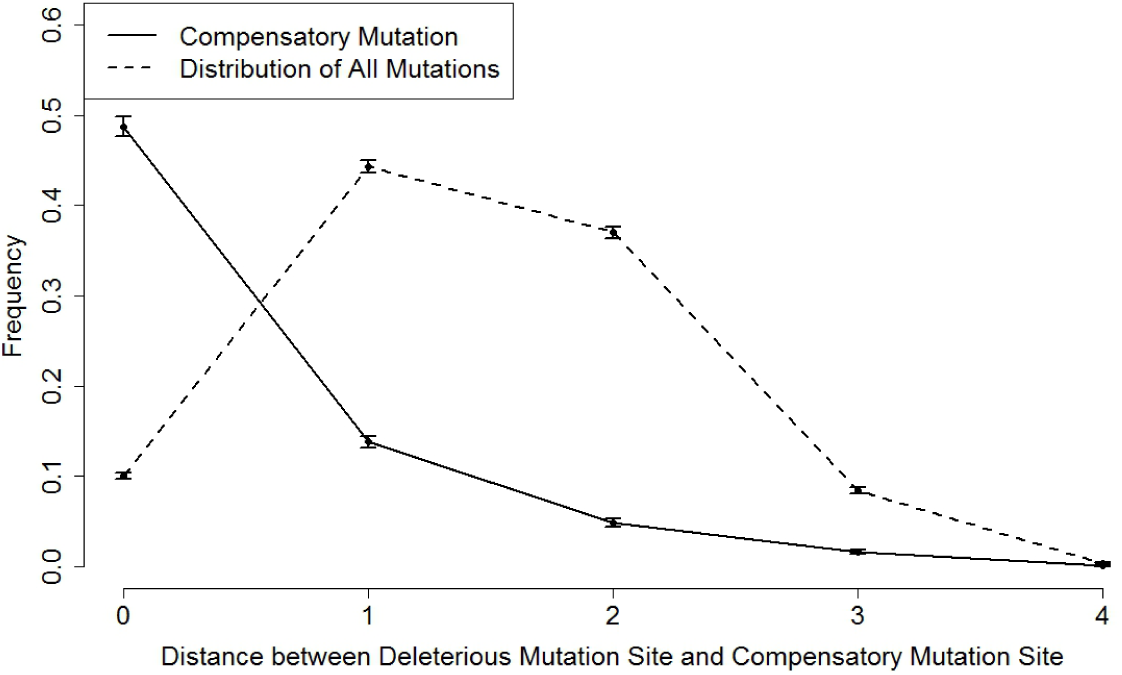
The compensatory mutation location and distance distribution of all mutations relative to the original deleterious mutation sites (Small Networks). For initially stable networks with size *N* = 5 and connectivity *c* = 0.4, we first collected a pool of compromised networks with deleterious mutations after a single mutation round. We then forced second mutations, classifying these as being 0 (on the same site), 1, 2, 3 and 4 steps away from the original deleterious mutations. For each of these mutation-site-distance categories, we measured the probability that the mutation was compensatory (that it returned the network to stability), based on 10, 000 sample networks collected for each distance category as shown in the solid line. We also recorded the spatial distribution of second mutations (10, 000 sample networks) occurring randomly in those compromised networks with respect to their original deleterious mutation sites, shown in the dashed line. The error bars represent 95% confidence intervals based on 100 independent runs.

**Figure S10:**
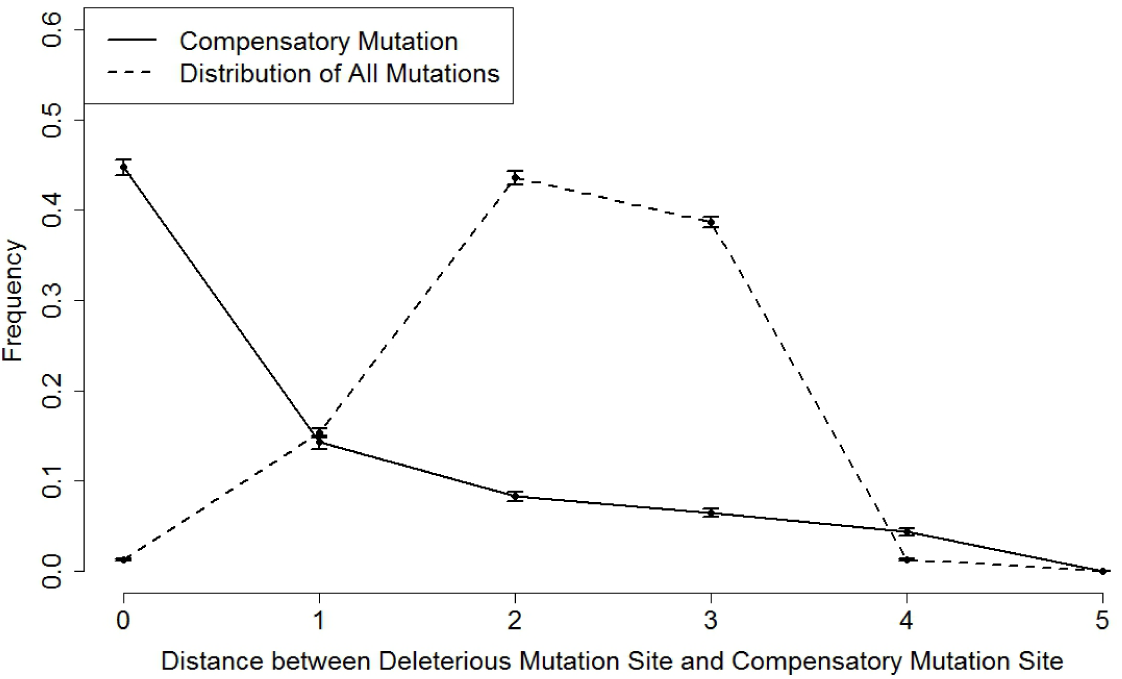
The compensatory mutation location and distance distribution of all mutations relative to the original deleterious mutation sites (Medium Networks). For initially stable networks with size *N* = 20 and connectivity *c* = 0.2, we first collected a pool of compromised networks with deleterious mutations after a single mutation round. We then forced second mutations, classifying these as being 0 (on the same site), 1, 2, 3, 4 and 5 steps away from the original deleterious mutations. For each of these mutation-site-distance categories, we measured the probability that the mutation was compensatory (that it returned the network to stability), based on 10, 000 sample networks collected for each distance category as shown in the solid line. We also recorded the spatial distribution of second mutations (10, 000 sample networks) occurring randomly in those compromised networks with respect to their original deleterious mutation sites, shown in the dashed line. The error bars represent 95% confidence intervals based on 100 independent runs.

**Figure S11:**
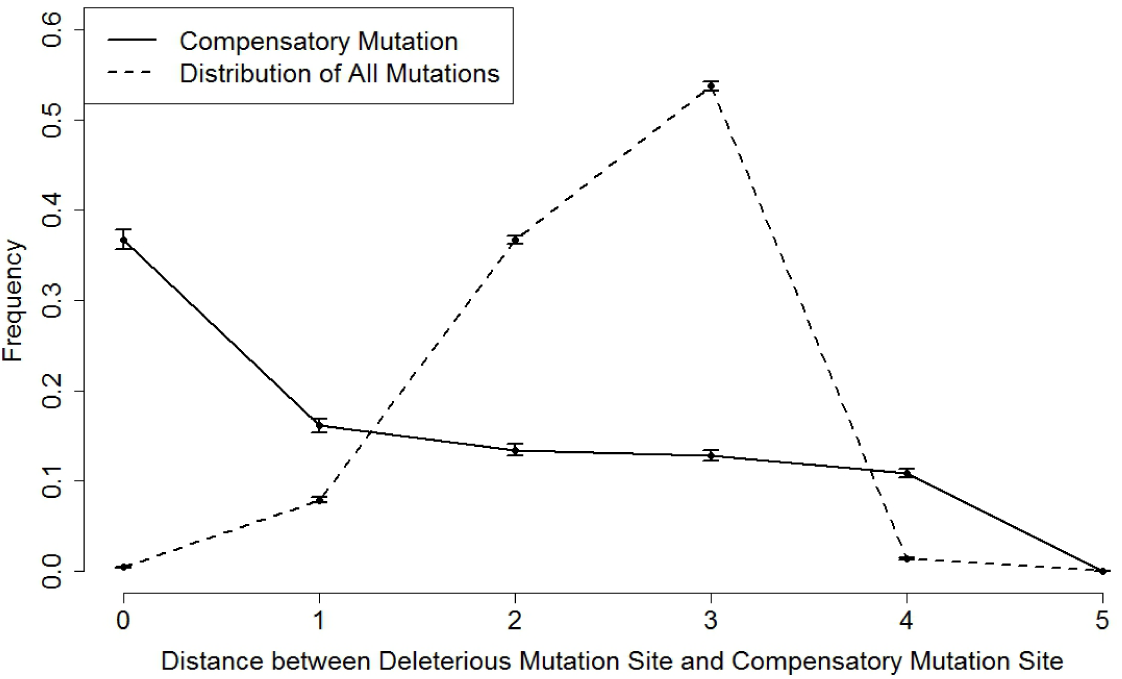
The compensatory mutation location and distance distribution of all mutations relative to the original deleterious mutation sites (Large Networks). For initially stable networks with size *N* = 40 and connectivity *c* = 0.15, we first collected a pool of compromised networks with deleterious mutations after a single mutation round. We then forced second mutations, classifying these as being 0 (on the same site), 1, 2, 3, 4 and 5 steps away from the original deleterious mutations. For each of these mutation-site-distance categories, we measured the probability that the mutation was compensatory (that it returned the network to stability), based on 10, 000 sample networks collected for each distance category as shown in the solid line. We also recorded the spatial distribution of second mutations (10, 000 sample networks) occurring randomly in those compromised networks with respect to their original deleterious mutation sites, shown in the dashed line. The error bars represent 95% confidence intervals based on 100 independent runs.

### Exploring the size of gene regulation on compensatory mutation frequency

In this set of experiments, we investigated effective changes in gene regulation associated with these mutations (see Fig. 3, main text, and also see Fig. S12). Specifically, we conducted experiments to measure the frequency of compensatory mutation when the second mutation had an additional weight added to it. We studied a range of weight changes from (*w* = [*−*5, 5]) with a step size of 0.05. For each step size, we first performed one mutation round as usual on the initial population of stable networks, creating a sub-population of 10, 000 compromised networks. Then, for these mutated networks we performed a second mutation round; however, this time instead of replacing one entry in the interaction matrix with *N* (0, 1), we added a fixed value *w* drawn from [*−*5, 5] to the original value of the randomly picked site. Then, we measured the frequency of second mutations restoring the network stability. We also performed similar experiments for medium (*N* = 20) and large networks (*N* = 40), as shown in Fig. S13 and Fig. S14.

**Figure S12:**
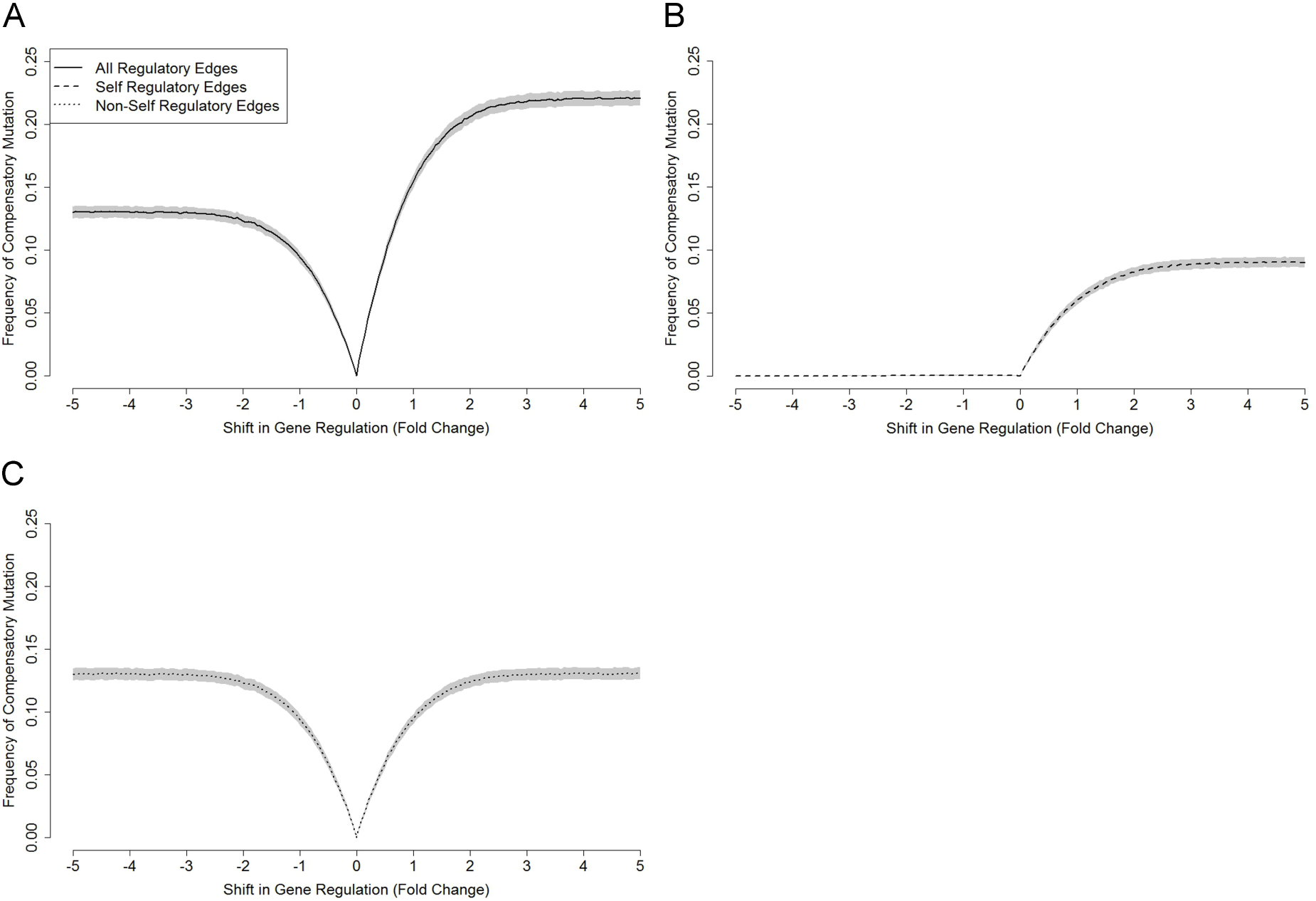
The influence of different intensities of gene regulations on the frequency of compensatory mutation (Small Networks). We first collected 10, 000 sample networks that had been made unstable by a single mutation from a pool of initially stable networks with *N* = 5 and *c* = 0.4. Then, we experimented with how a new mutation of varying intensities of gene regulation altered the chances of restoring gene stability. Specifically, we performed new mutations to those compromised networks with deleterious mutations by adding a weight from [−5, +5] (step size 0.5) to the original regulatory impact, then assessed the resulting patterns in all regulatory edges (**A**), in self-regulatory edges (**B**) and ignoring self-regulatory edges (**C**). The shaded areas represent 95% confidence intervals based on 100 independent runs.

**Figure S13:**
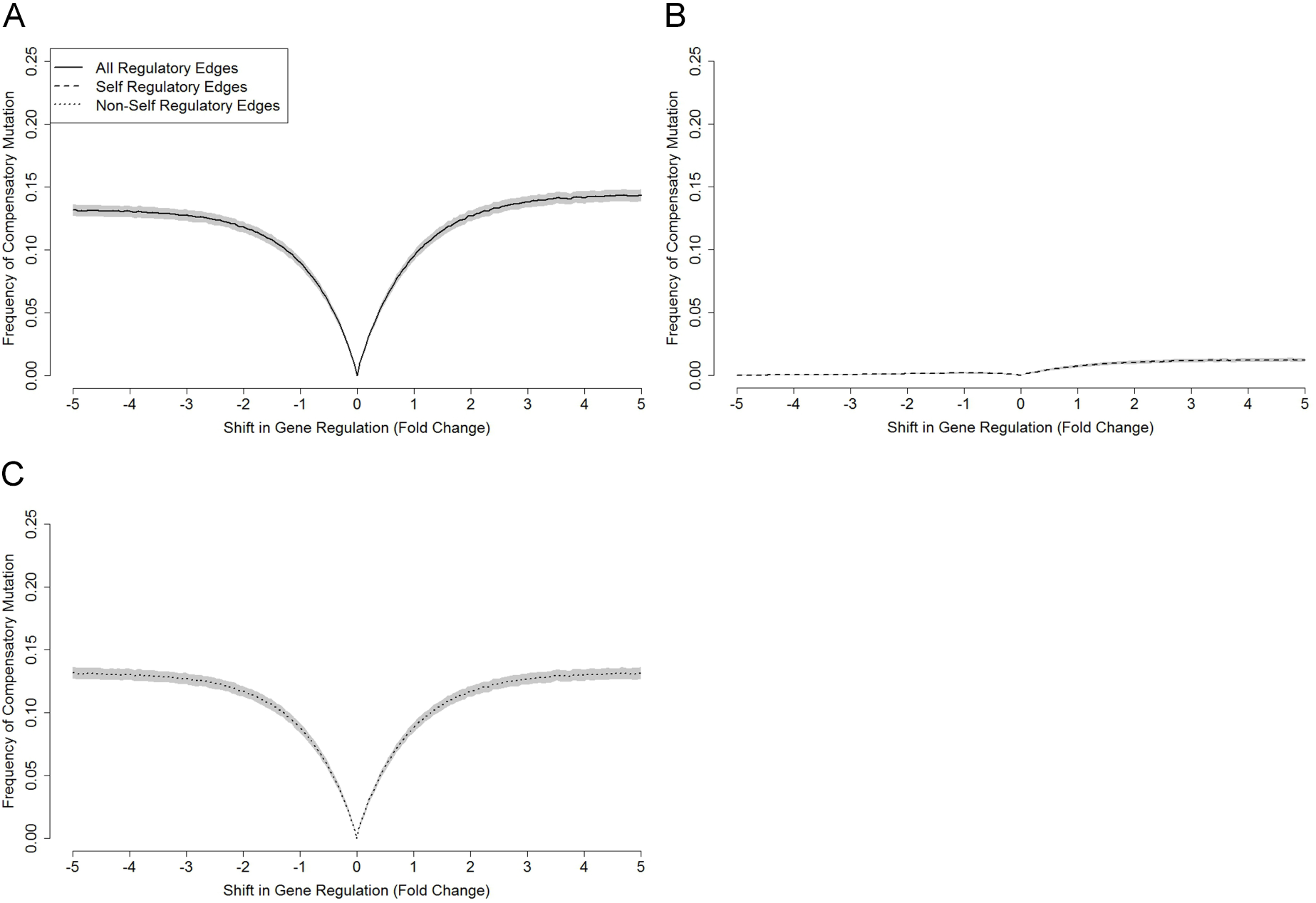
The influence of different intensities of gene regulation on frequency of compensatory mutation (Medium Networks). We first collected 10, 000 sample networks that had been made unstable by a single mutation from a pool of initially stable networks with *N* = 20 and *c* = 0.2. Then, we experimented with how a new mutation of varying intensities of gene regulation altered the chances of restoring gene stability. Specifically, we performed new mutations to those compromised networks with deleterious mutations by adding a weight from [−5, +5] (step size 0.5) to the original regulatory impact, then assessed the resulting patterns in all regulatory edges (**A**), in self-regulatory edges (**B**) and ignoring self-regulatory edges (**C**). The shaded areas represent 95% confidence intervals based on 100 independent runs.

**Figure S14:**
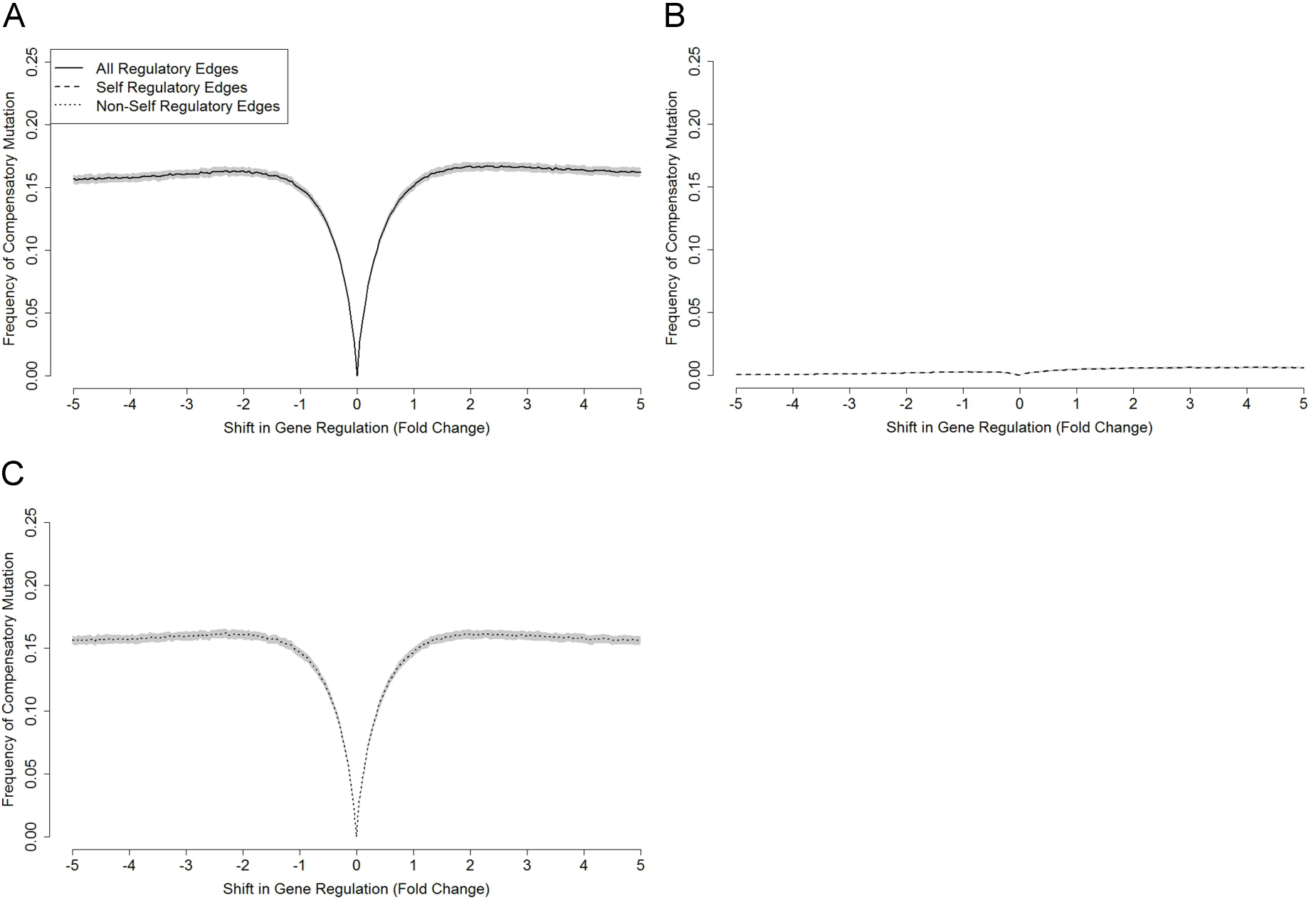
The influence of different intensities of gene regulation on frequency of compensatory mutation (Large Networks). We first collected 10, 000 sample networks that had been made unstable by a single mutation from a pool of initially stable networks with *N* = 40 and *c* = 0.15. Then, we experimented with how a new mutation of varying intensities of gene regulation altered the chances of restoring gene stability. Specifically, we performed new mutations to those compromised networks with deleterious mutations by adding a weight from [−5, +5] (step size 0.5) to the original regulatory impact, then assessed the resulting patterns in all regulatory edges (**A**), in self-regulatory edges (**B**) and ignoring self-regulatory edges (**C**). The shaded areas represent 95% confidence intervals based on 100 independent runs.

### Exploring the distribution of regulation in initially-stable, compromised and restored networks

In this set of experiments, we investigated the distribution of regulation in initially stable, compromised and restored networks (see Fig. S5). Specifically, we collected 10, 000 sample regulatory values each from edges of randomly generated stable networks, edges where deleterious mutations occurred (compromising network stability), and edges where compensatory mutations occurred (restoring previously compromised networks). We then measured their corresponding distributions, discriminating between self- and non-self-regulatory edges. We also performed similar experiments for medium (*N* = 20) and large networks (*N* = 40), as shown in Fig. S15 and Fig. S16.

**Figure S15:**
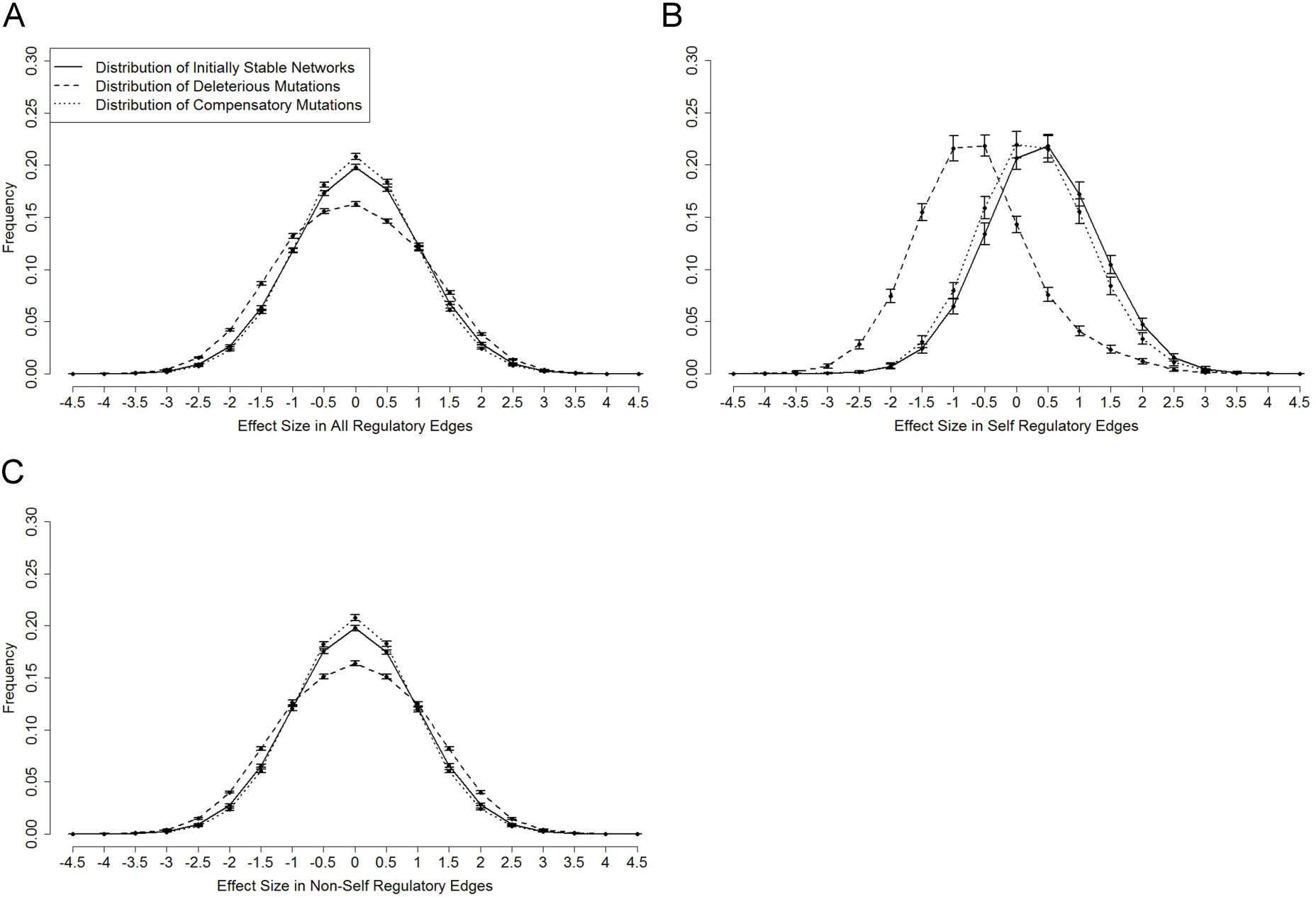
The distribution of regulation in initially stable, compromised and restored networks (Medium Networks). For randomly generated stable networks with *N* = 20 and *c* = 0.2, we collected 10, 000 sample regulations. We also collected 10, 000 sample regulation weights from deleterious mutations that compromised initially stable networks as well as from compensatory mutations that restored the stability of previously broken networks. We then measured the distributions in all regulatory edges (**A**), in self-regulatory edges (**B**) and ignoring self-regulatory edges (**C**). Given that the regulations are continuous values, we grouped them into 19 bins from [−4.5, +4.5] (step size 0.5). The error bars represent 95% confidence intervals based on 100 independent runs.

**Figure S16:**
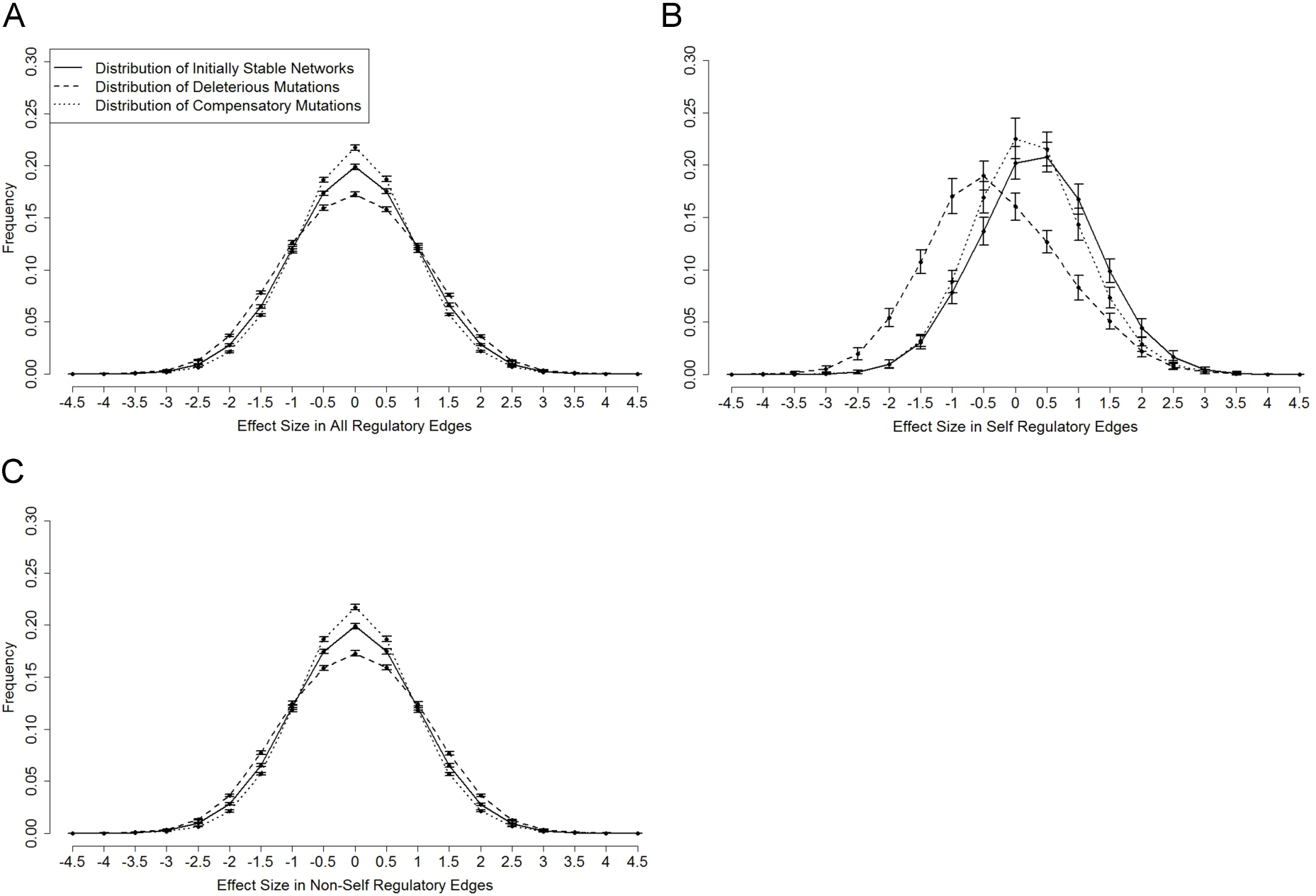
The distribution of regulation in initially stable, compromised and restored networks (Large Networks). For randomly generated stable networks with *N* = 40 and *c* = 0.15, we collected 10, 000 sample regulations. We also collected 10, 000 sample regulation weights from deleterious mutations that compromised initially stable networks as well as from compensatory mutations that restored the stability of previously broken networks. We then measured the distributions in all regulatory edges (**A**), in self-regulatory edges (**B**) and ignoring self-regulatory edges (**C**). Given that the regulations are continuous values, we grouped them into 19 bins from [−4.5, +4.5] (step size 0.5). The error bars represent 95% confidence intervals based on 100 independent runs.

### Exploring properties of location and size effects in neutral mutations

In this set of experiments, we investigated properties of location and size effects in neutral mutations which served as control groups for solid lines in Fig. 3A and B, main text, Specifically, to test the location effect, we collected a population pool of stable networks that had been subjected to one round of mutation (neutral). Then, we measured the probability of stable networks after performing a second mutation that was 0, 1, 2, 3, and 4 steps away from the previous neutral mutation site based on 10, 000 sample networks for each distance category (see dashed line in Fig. 3A). Similarly, to test the mutation size effect, we collected a population pool of stable networks that had been subjected to one round of mutation (neutral). Then, we measured the probability of stable networks after performing a second mutation that had a particular shift in gene regulation from [*−*5, +5] based on 10, 000 sample networks for each shifted-weight category (see dashed line in Fig. 3B). In both tests for location and size effects, we also performed similar experiments for medium (*N* = 20) and large networks (*N* = 40), as shown in Fig. S17 and Fig. S18.

**Figure S17:**
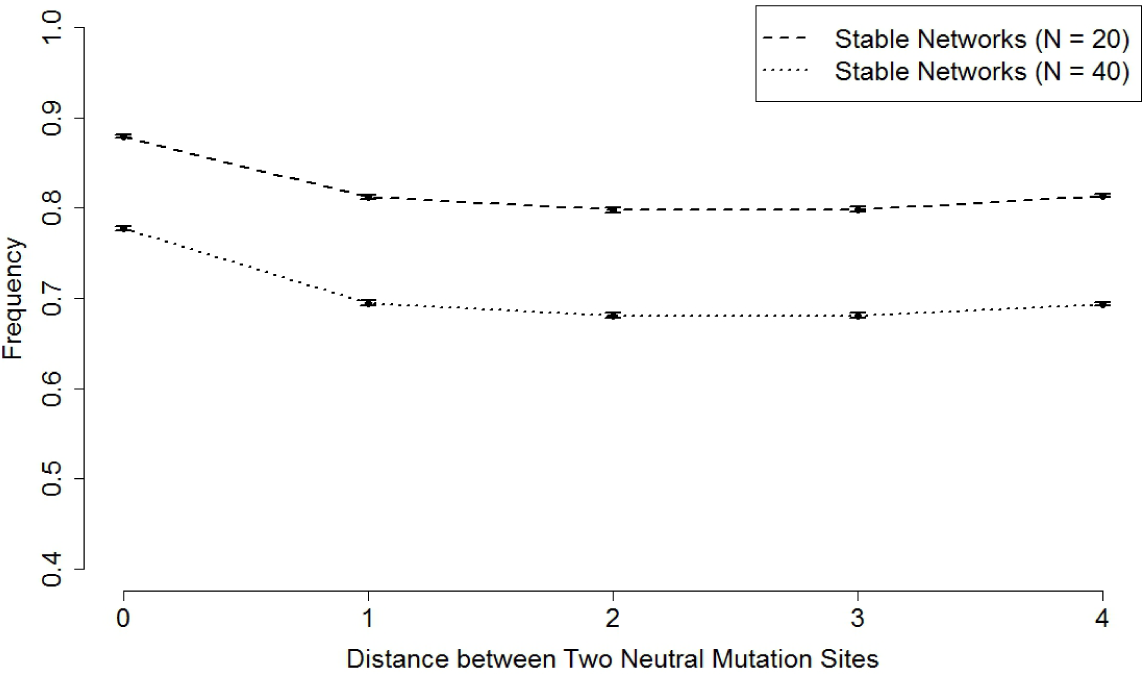
Location effect in networks with neutral mutations (Medium and Large Networks). For medium networks (*N* = 20*, c* = 0.2) and large networks (*N* = 40*, c* = 0.15), we first collected a pool of stable networks with neutral mutations after a single mutation round. We then forced second mutations, classifying these as being 0 (on the same site), 1, 2, 3 and 4 steps away from the previous neutral mutations. For each of these mutation-site-distance categories, we measured the probability that the mutation was neutral (did not impair network stability) based on 10, 000 sample networks collected for each distance category. The error bars represent 95% confidence intervals based on 100 independent runs.

**Figure S18:**
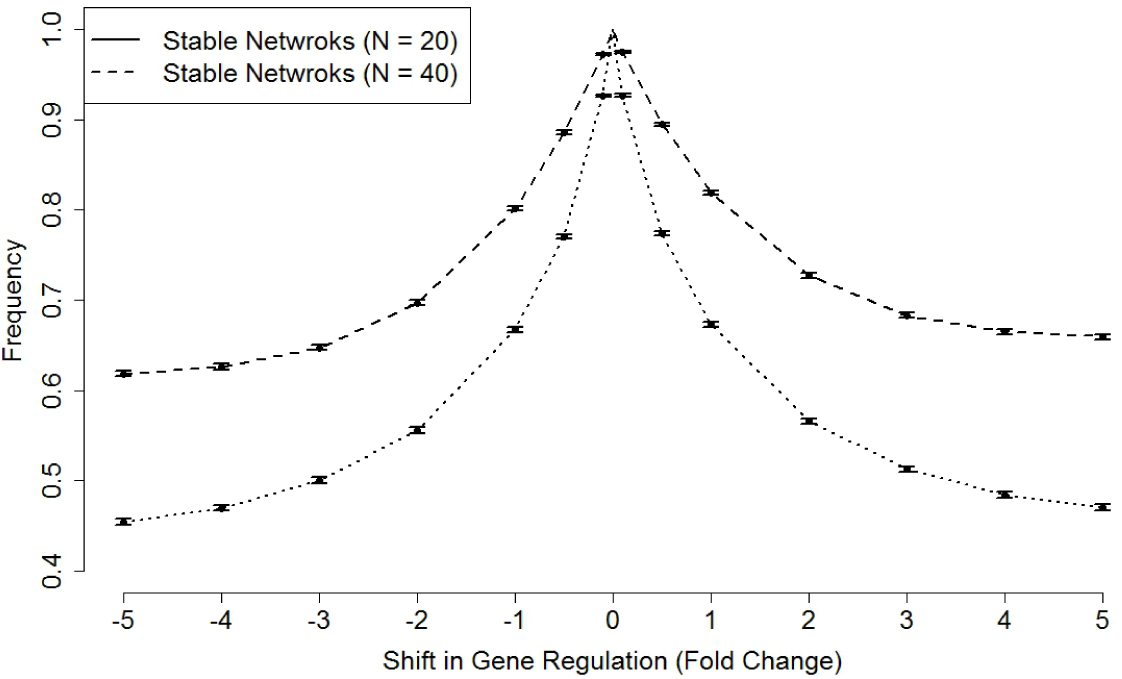
Mutation size effect in networks with neutral mutations (Medium and Large Networks). We first collected 10, 000 stable networks with neutral mutations after a single mutation round from a pool of initially stable medium networks (*N* = 20*, c* = 0.2) and large networks (*N* = 40*, c* = 0.15). Then, we experimented with how new mutations of varying intensities of gene regulation altered the chance of retaining network stability. Specifically, we performed new mutations to those networks with neutral mutations by adding a weight from [−5, +5] (step size 1 and with four additional regulation shifts: 0.5, 0.1, 0.1 and 0.5) to the original regulatory impact, then assessed the resulting patterns. The error bars represent 95% confidence intervals based on 100 independent runs.

### Exploring the impact of distance and size effects on network robustness

In this set of experiments, we explored the effects of location and mutation size on robustness in networks with one deleterious mutation and one compensatory mutation and in networks with two consecutive neutral mutations to investigate whether networks with compensatory mutations have a different evolutionary consequence compare with networks with neutral mutations (see Fig. 3C and D and also see Fig. S19 and Fig. S22).

Specifically, to test the distance effect, we collected 10, 000 sample networks at each distance (between deleterious mutation and compensatory mutation). Then, for each category of distance, we measured the proportion of stable networks after one additional round of single mutation. The reported results are both actual robustness (see the solid line in Fig. S19A) and percentage change in robustness (see the solid line in Fig. S19B). Similarly, for the control group, instead of collecting networks that were subjected to one deleterious mutation and one subsequent compensatory mutation, we collected 10, 000 sample networks that were subjected to two consecutive neutral mutations at each distance (between two neutral mutations), and then assessed the actual robustness (see the dashed line in Fig. S19A) as well as the percentage of robustness change (see the dashed line in Fig. S19B). We also performed similar experiments for medium (*N* = 20) and large networks (*N* = 40), as shown in Fig. S20 and Fig. S21.

Likewise, to test size effect, we collected 10, 000 sample networks that were compensated by mutations with different shifts in gene regulation. Then, for each category of mutation size, we measured the proportion of stable networks after one additional round of single mutation. The reported results are both actual robustness (see the solid line in Fig. S22A) and percentage change in robustness (see the solid line in Fig. S22B). Similarly, for the control group, instead of collecting networks that were subjected to one normal deleterious mutation and one subsequent compensatory mutation with different shifts in gene regulation, we collected 10, 000 sample networks that were subjected to two consecutive neutral mutations, one normal neutral mutation and the other neutral mutation with different shifts in gene regulation, and then assessed the actual robustness (see the dashed line in Fig. S22A) as well as the percentage of robustness change (see the dashed line in Fig. S22B). We also performed similar experiments for medium (*N* = 20) and large networks (*N* = 40), as shown in Fig. S23 and Fig. S24.

**Figure S19:**
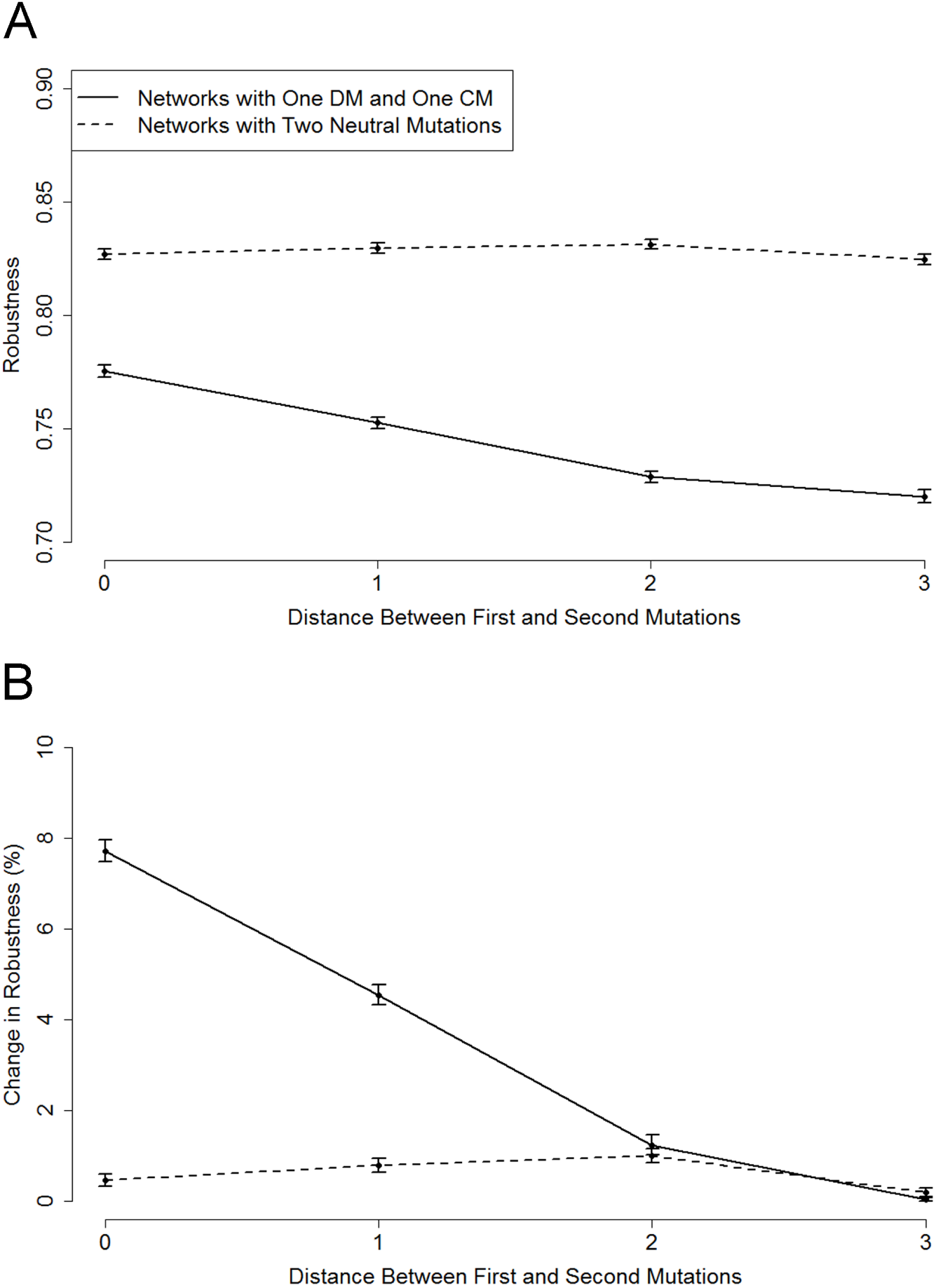
The impact of distance effect on network robustness (Small Networks). For small networks (*N* = 5*, c* = 0.4), we collected 10, 000 sample stable networks that were subjected one deleterious mutation and then restored by one subsequent compensatory mutation that was 0, 1, 2 and 3 steps away from the previous deleterious mutation. The sample networks for control group were collected in a similar way, except that the networks were subjected to two consecutive neutral mutations. Then, we assessed robustness of sample networks at each distance step. The reported results are actual robustness (**A**), and change in robustness (**B**) (the actual robustness was normalised by subtracting the minimal value among all categories, and then divided by the minimal value). The error bars represent 95% confidence intervals based on 100 independent runs.

**Figure S20:**
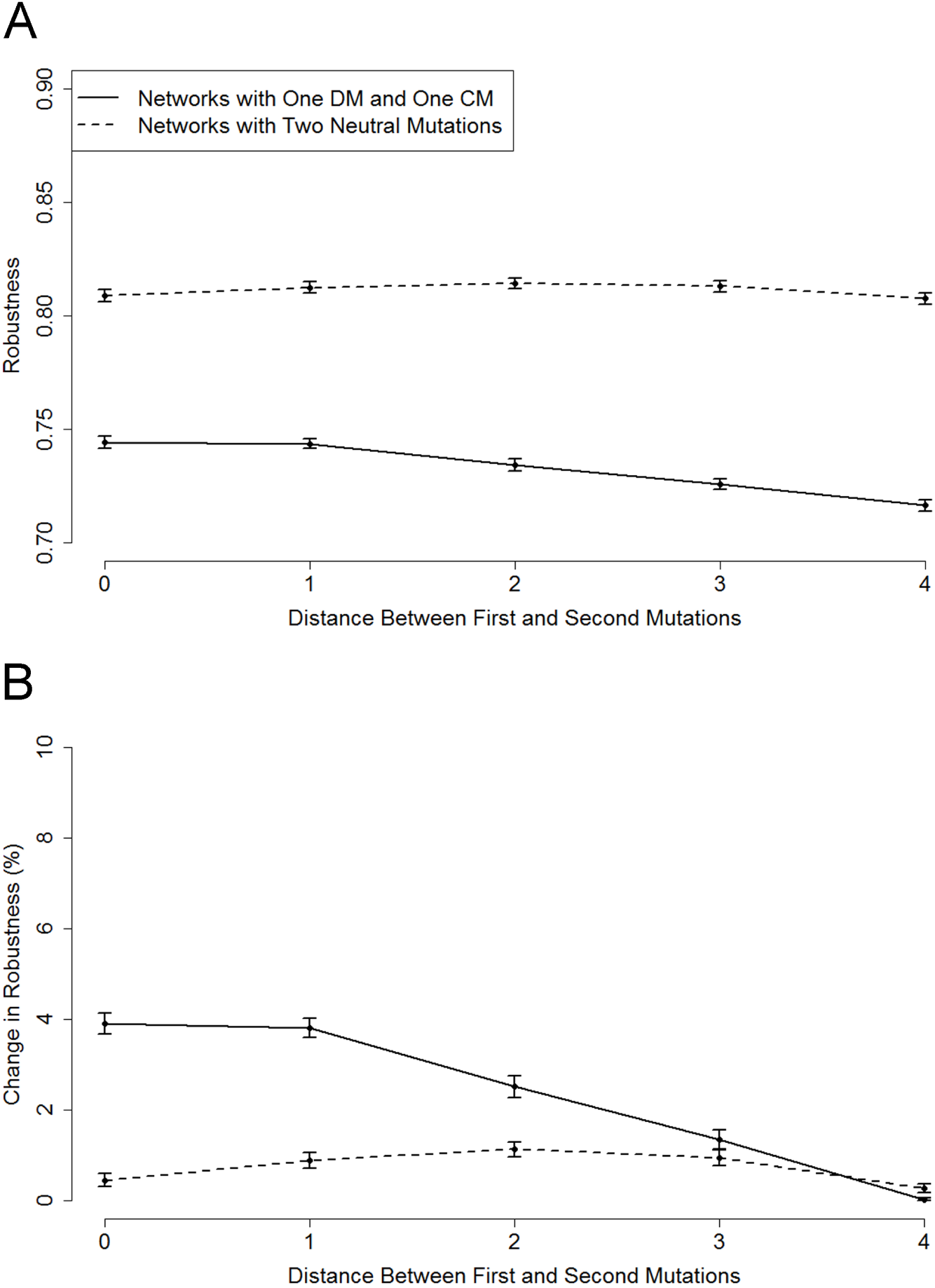
The impact of distance effect on network robustness (Medium Networks). For medium networks (*N* = 20*, c* = 0.2), we collected 10, 000 sample stable networks that were subjected one deleterious mutation and then restored by one subsequent compensatory mutation that was 0, 1, 2, 3 and 4 steps away from the previous deleterious mutation. The sample networks for the control group were collected in a similar way, except that the networks were subjected to two consecutive neutral mutations. Then, we assessed the robustness of the sample networks at each distance step. The reported results are actual robustness (**A**), and change in robustness (**B**) (the actual robustness was normalised by subtracting the minimal value among all categories, and then dividing by the minimal value). The error bars represent 95% confidence intervals based on 100 independent runs.

**Figure S21:**
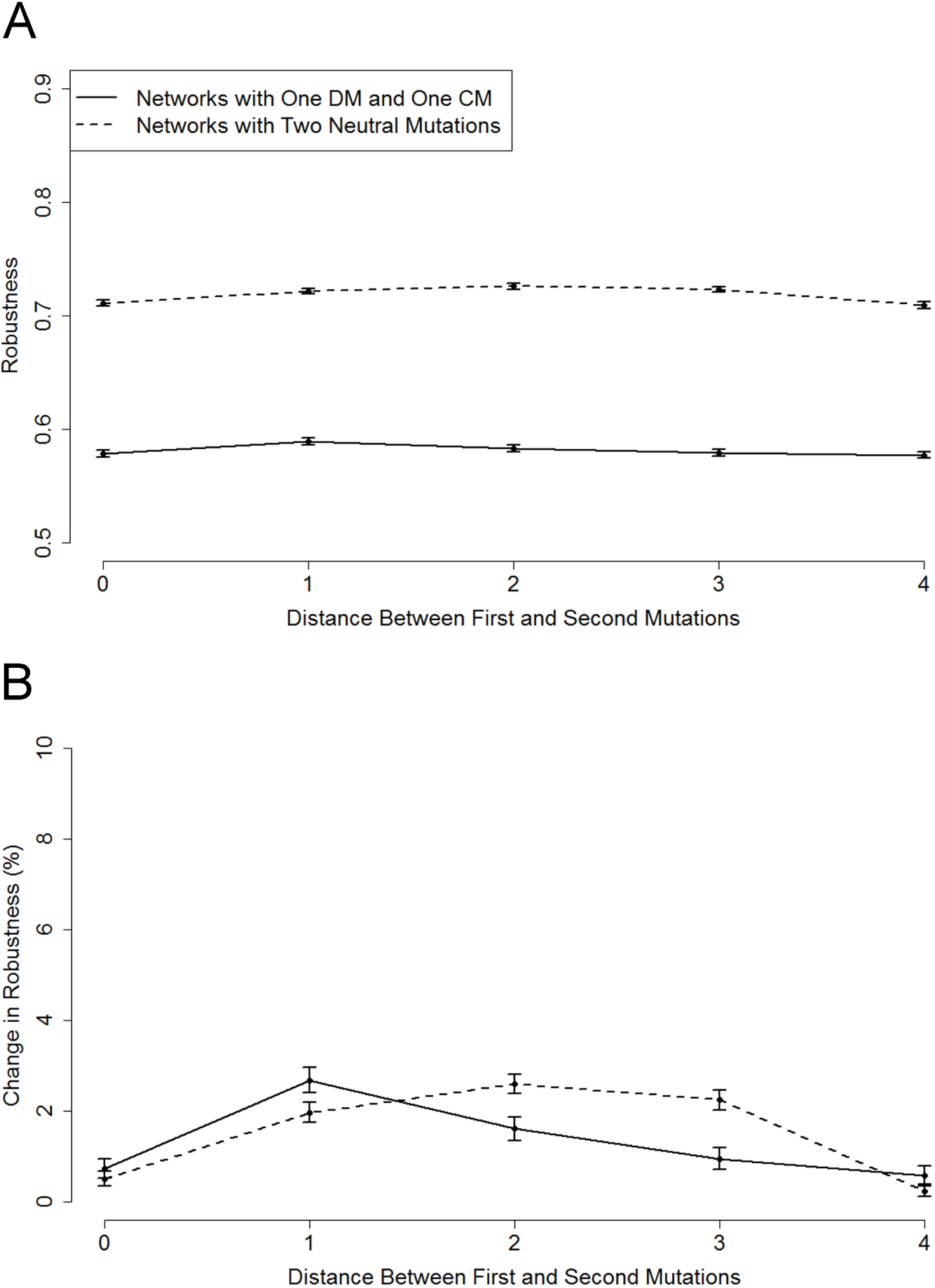
The impact of distance effect on network robustness (Large Networks). For large networks (*N* = 40*, c* = 0.15), we collected 10, 000 sample stable networks that were subjected one deleterious mutation and then restored by one subsequent compensatory mutation that was 0, 1, 2, 3 and 4 steps away from the previous deleterious mutation. The sample networks for the control group were collected in a similar way, except that the networks were subjected to two consecutive neutral mutations. Then, we assessed the robustness of the sample networks at each distance step. The reported results are actual robustness (**A**), and change in robustness (**B**) (the actual robustness was normalised by subtracting the minimal value among all categories, and then dividing by the minimal value). The error bars represent 95% confidence intervals based on 100 independent runs.

**Figure S22:**
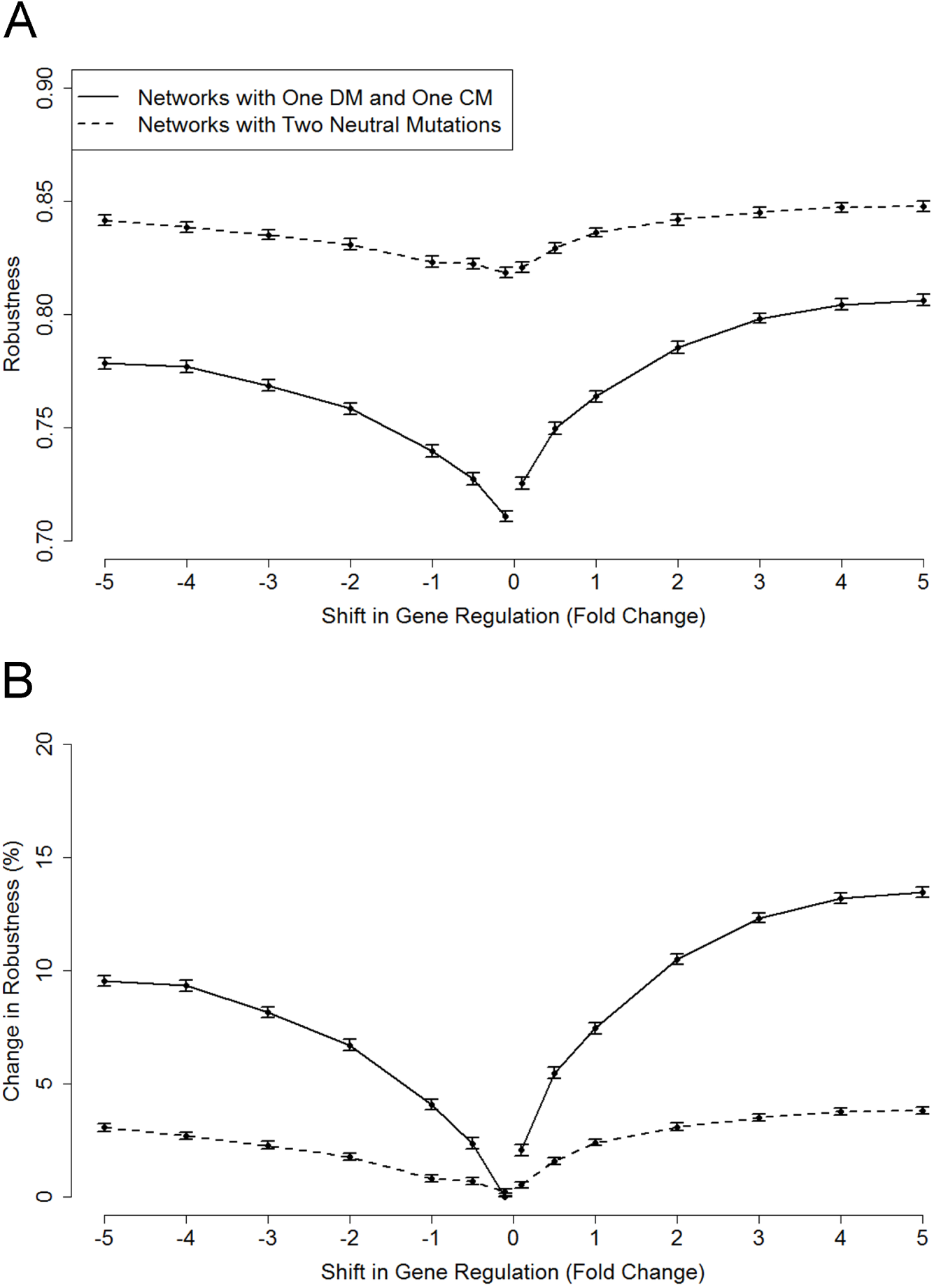
The impact of mutation size effect on network robustness (Small Networks). For small networks (*N* = 5*, c* = 0.4), we collected 10, 000 sample stable networks that were subjected one deleterious mutation and then restored by one subsequent compensatory mutation with different shifts in gene regulation from [−5, +5] (step size 1 and with four additional regulation shifts: 0.5, 0.1, 0.1 and 0.5). The sample networks for control group were collected in a similar way, except that the networks were subjected to two consecutive neutral mutations. Note that the second neutral mutation has different shifts in gene regulation as the compensatory mutation. Then, we assessed robustness of sample networks at each category. The reported results are actual robustness (**A**), and change in robustness (**B**) (the actual robustness was normalised by subtracting the minimal value among all categories, and then divided by the minimal value). The error bars represent 95% confidence intervals based on 100 independent runs.

**Figure S23:**
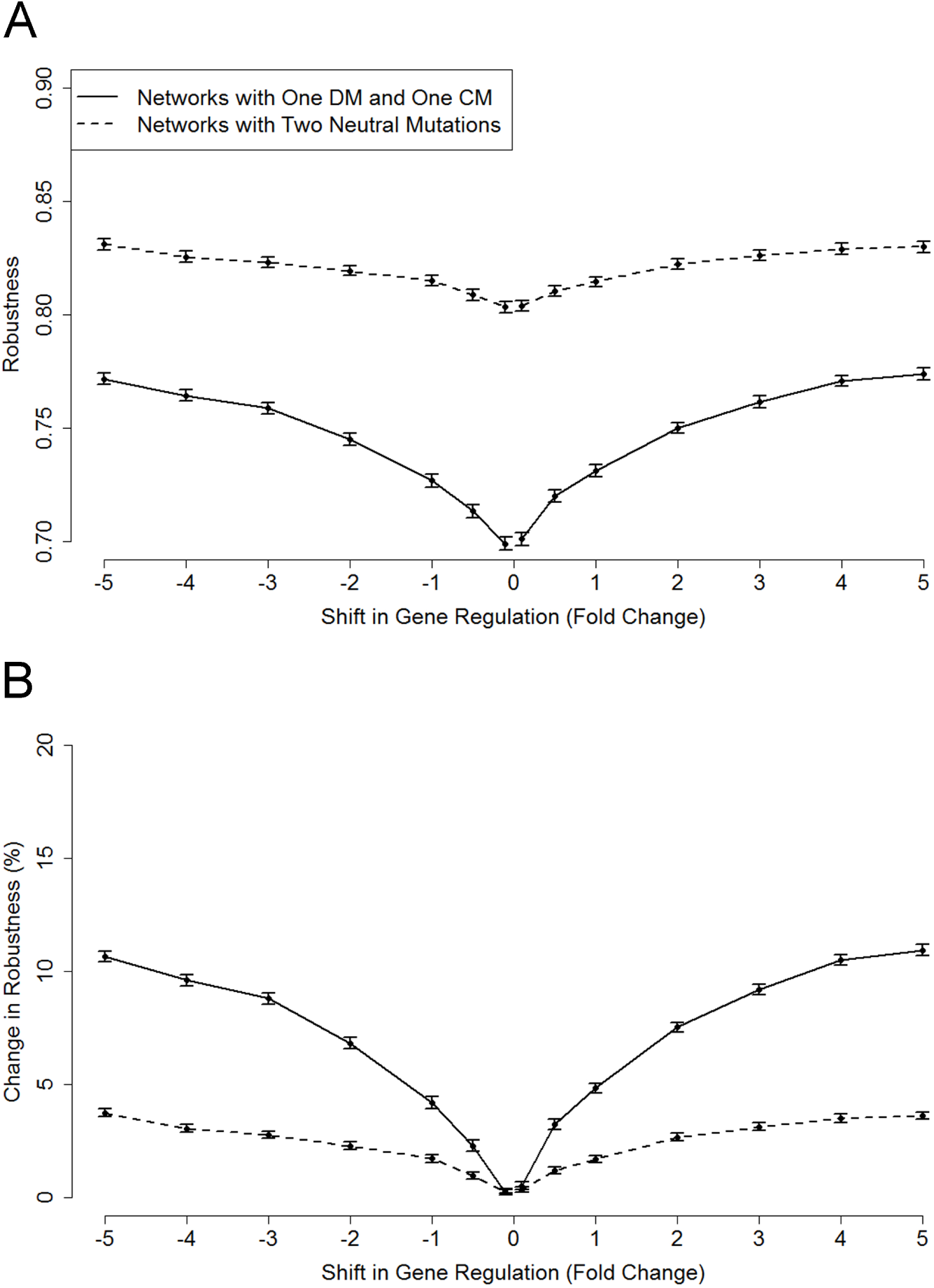
The impact of mutation size effect on network robustness (Medium Networks). For medium networks (*N* = 20*, c* = 0.2), we collected 10, 000 sample stable networks that were subjected one deleterious mutation and then restored by one subsequent compensatory mutation with different shifts in gene regulation from [−5, +5] (step size 1 and with four additional regulation shifts: 0.5, 0.1, 0.1 and 0.5). The sample networks for the control group were collected in a similar way, except that the networks were subjected to two consecutive neutral mutations. Note that the second neutral mutation has different shifts in gene regulation to the compensatory mutation. Then, we assessed the robustness of the sample networks at each category. The reported results are actual robustness (**A**), and change in robustness (**B**) (the actual robustness was normalised by subtracting the minimal value among all categories, and then dividing by the minimal value). The error bars represent 95% confidence intervals based on 100 independent runs.

**Figure S24:**
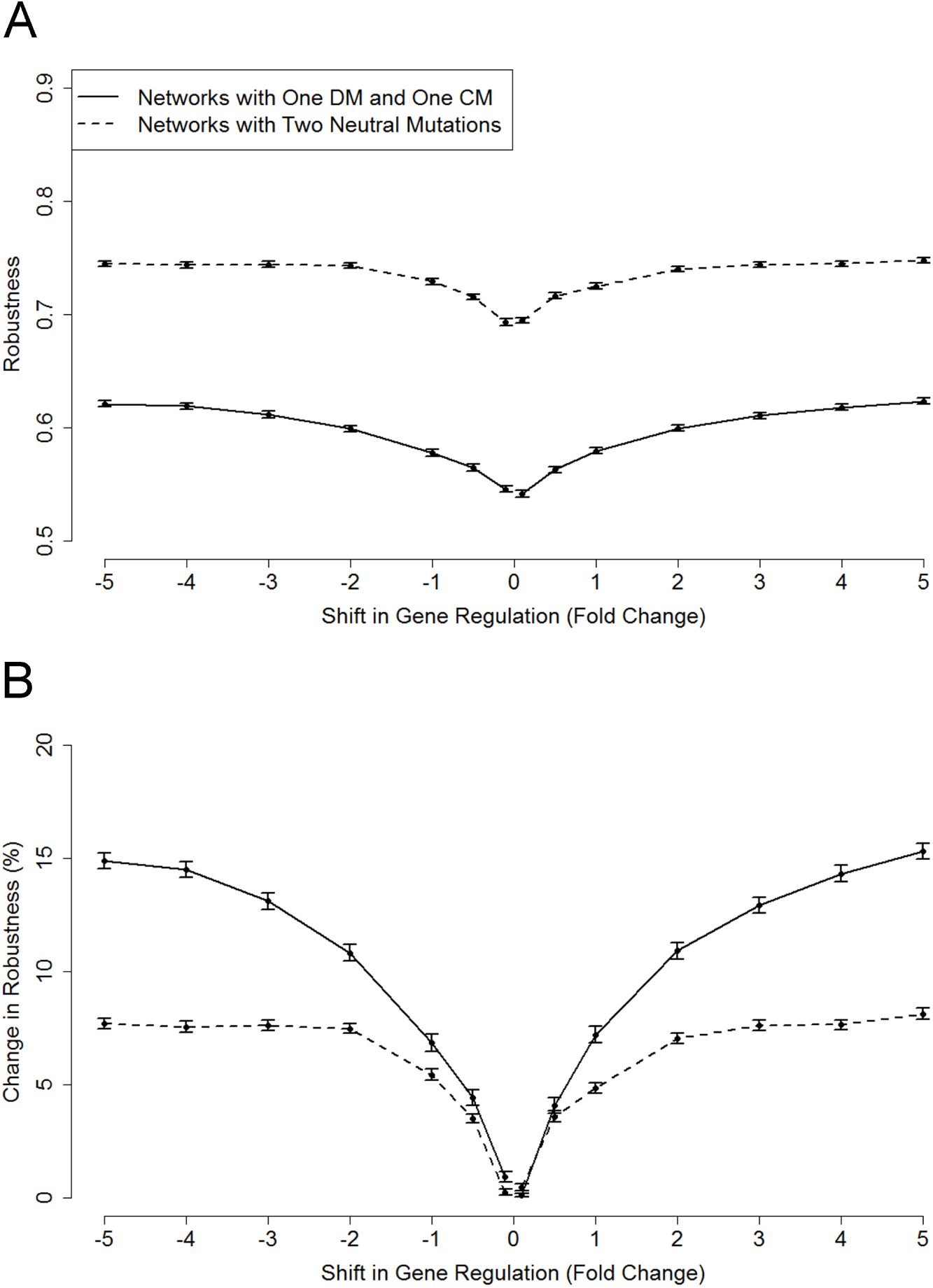
The impact of mutation size effect on network robustness (Large Networks). For large networks (*N* = 40*, c* = 0.15), we collected 10, 000 sample stable networks that were subjected one deleterious mutation and then restored by one subsequent compensatory mutation with different shifts in gene regulation from [−5, +5] (step size 1 and with four additional regulation shifts: 0.5, 0.1, 0.1 and 0.5). The sample networks for the control group were collected in a similar way, except that the networks were subjected to two consecutive neutral mutations. Note that the second neutral mutation has different shifts in gene regulation to the compensatory mutation. Then, we assessed the robustness of the sample networks at each category. The reported results are actual robustness (**A**), and change in robustness (**B**) (the actual robustness was normalised by subtracting the minimal value among all categories, and then dividing by the minimal value). The error bars represent 95% confidence intervals based on 100 independent runs.

### Exploring how network connectivity evolves under a relaxed selection regime

In this set of experiments, we investigated whether regulatory complexity (increased network connectivity) could arise under a relaxed selection regime where compensatory mutations could occur and accumulate (see Fig. 4 and Fig. S6).

In the first set of experiments, we tested whether we could observe greater complexity arising using a population pool of 10, 000 stable networks of *N* = 10 genes with a simple ‘Star’ topology (see fig 4, main text). Specifically, the initial population pool was generated using the following rules:

- Randomly select a gene to be the hub node.
- There is at least one edge between the hub node and non-hub nodes (either inward or outward); there is a possibility (0.5) of having both inward and outward edges.
- Each node has a possibility (0.5) of having a self-regulatory edge (including the hub node).
- The value (interaction strength) of each edge is drawn from the standard normal distribution *N* (0, 1).

In theory, for network size *N* = 10, the minimum connectivity is *c*_min_ = 0.09 (9 edges) and the maximum connectivity is *c*_max_ = 0.28 (28 edges). In the randomly generated initial population pool used in this paper, the minimum connectivity was *c*_min_ = 0.10 (10 edges), the maximum connectivity was *c*_max_ = 0.26 (26 edges), the median connectivity was *c͂* = 0.17 (17 edges) and the average connectivity was *c̄ ≈* 0.17. Then, the initial population was evolved for 5, 000 generations under strong and relaxed selection regimes: In four scenarios with selection for network stability, the initial population was evolved under: a no mutation and no recombination regime, a mutation but no recombination regime, a recombination but no mutation regime, a mutation and recombination; in three other scenarios, the initial population was evolved under a relaxed selection regime with a frequency of 1*/*10, 1*/*25, and 1*/*50. The statistical details for connectivity in initial and evolved populations can be found in Table S1. Note that compensatory mutation could only occur during periods of relaxed selection.

In order to further test the hypothesis that relaxed selection can facilitate regulatory complexity, in the second set of experiments, we further investigated how network connectivity evolves under a relaxed selection regime using randomly generated networks (see Fig. S6). Specifically, for a network size *N* = 40 with connectivity *c* = 0.15, we collected 10, 000 stable networks, each of which had the same initial gene expression pattern, all activation, i.e., **s**(0) = (+1, +1, …, +1). This population was then evolved for 5, 000 generations, in this case allowing for recombination with other individuals from the same generation. Note that in the previously-described experiments in this paper, a mutation could not change the topology of an individual network; that is, it could not change zero elements into non-zero or *vice versa*. In contrast, recombination can alter the topology if the non-zero sites are different in individual networks. The reported results are the mean network connectivity of all individuals in the population in every 200 generations under different frequencies of relaxed selection. Note that network connectivity was measured in the next generation of network stability selection immediately after the previous relaxed selection; therefore, we only report the results in stable networks.

**Table S1:**
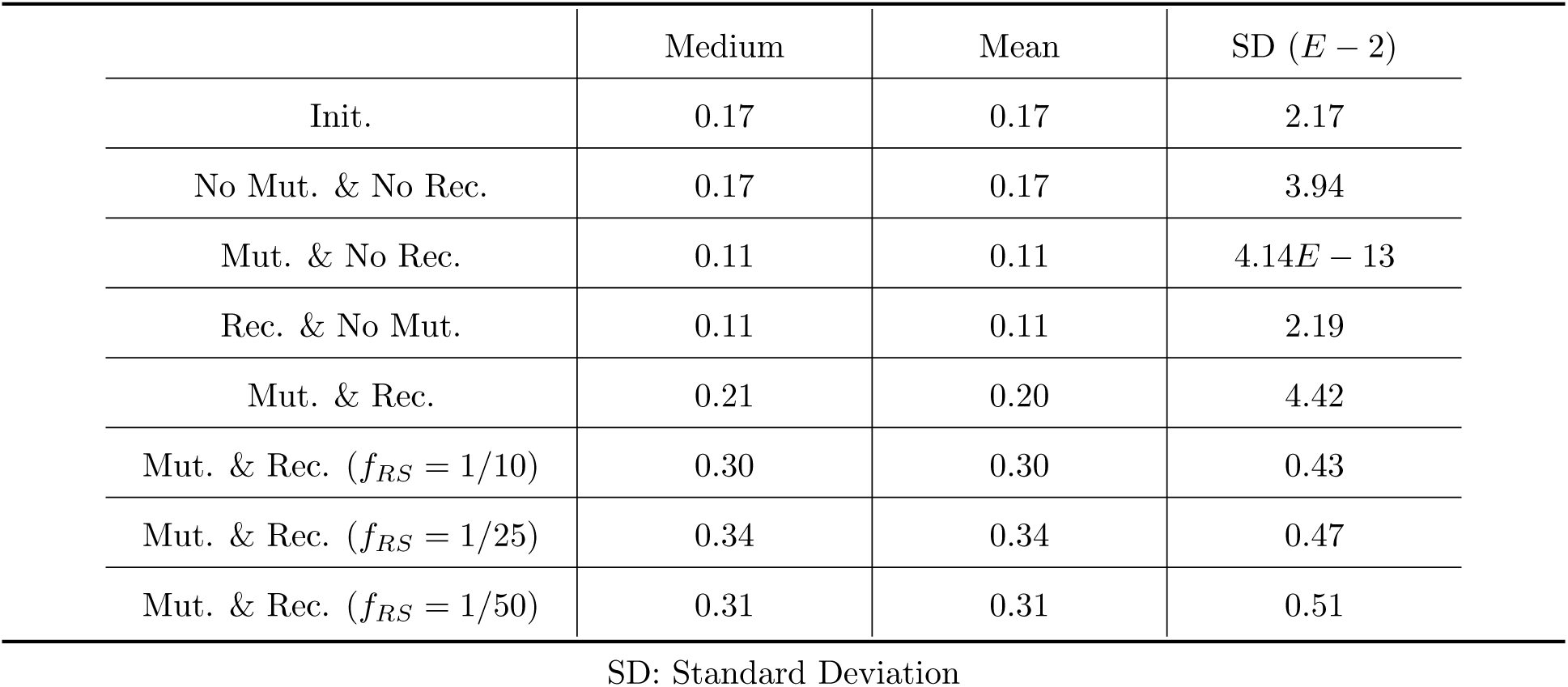
Basic statistics of evolved networks with a ‘Star’ topology

## Footnotes

1 Note that robustness is higher overall for networks having experienced two neutral mutations—this is unlikely to be caused by the mutations, and more likely to be a characteristic of the network likely to contain neutral mutations.

## References

C. O. Wilke and C. Adami. Interaction between directional epistasis and average mutational effects. Proceedings of the Royal Society of London B: Biological Sciences, 268(1475):1469–1474, 2001.

Claus Wilke, Richard Lenski, and Christoph Adami. Compensatory mutations cause excess of antagonistic epistasis in RNA secondary structure folding. BMC Evolutionary Biology, 3(1):3, 2003.

Niko Beerenwinkel, Lior Pachter, Bernd Sturmfels, Santiago Elena, and Richard Lenski. Analysis of epistatic interactions and fitness landscapes using a new geometric approach. BMC Evolutionary Biology, 7(1):60, 2007.

B. Lehner. Molecular mechanisms of epistasis within and between genes. Trends in Genetics, 27(8):323–331, 2011.

D. R. Rokyta, P. Joyce, S. B. Caudle, C. Miller, C. J. Beisel, and H. A. Wichman. Epistasis between beneficial mutations and the phenotype-to-fitness map for a ssDNA virus. PLoS Genetics, 7(6):e1002075, 2011.

Solip Park and Ben Lehner. Epigenetic epistatic interactions constrain the evolution of gene expression. Molecular Systems Biology, 9:645–645, 2013.

S. Ciliberti, O. C. Martin, and A. Wagner. Innovation and robustness in complex regulatory gene networks. Proceedings of the National Academy of Sciences of the United States of America, 104(34):13591–13596, 2007.

Anton Crombach and Paulien Hogeweg. Evolution of evolvability in gene regulatory networks. PLoS Computational Biology, 4(7):e1000112, 2008.

Philip A. Romero and Frances H. Arnold. Exploring protein fitness landscapes by directed evolution. Nature Reviews Molecular Cell Biology, 10(12):866–876, 2009.

Masaki E. Tsuda and Masakado Kawata. Evolution of gene regulatory networks by fluctuating selection and intrinsic constraints. PLoS Computational Biology, 6(8):e1000873, 2010.

Carrie F. Olson-Manning, Maggie R. Wagner, and Thomas Mitchell-Olds. Adaptive evolution: Evaluating empirical support for theoretical predictions. Nature Reviews Genetics, 13(12):867–877, 2012.

James Cotterell and James Sharpe. Mechanistic explanations for restricted evolutionary paths that emerge from gene regulatory networks. PLoS ONE, 8(4):e61178, 2013.

Andreas Wagner and Jeremiah Wright. Alternative routes and mutational robustness in complex regulatory networks. Biosystems, 88(1-2):163–172, 2007.

Michael Lynch, Matthew S Ackerman, Jean-Francois Gout, Hongan Long, Way Sung, W Kelley Thomas, and Patricia L Foster. Genetic drift, selection and the evolution of the mutation rate. Nature Reviews Genetics, 17(11):704–714, 2016.

Adam M Siepielski, Joseph D DiBattista, and Stephanie M Carlson. It’s about time: the temporal dynamics of phenotypic selection in the wild. Ecology Letters, 12(11):1261–1276, November 2009a.

Anett Dunai, Réka Spohn, Zoltán Farkas, Viktória Lázár, Ádám Györkei, Gábor Apjok, Gábor Boross, Balázs Szappanos, Gábor Grézal, Anikó Faragó, László Bodai, Balázs Papp, and Csaba Pál. Rapid decline of bacterial drug-resistance in an antibiotic-free environment through phenotypic reversion. eLife, 8:e47088, aug 2019. ISSN 2050-084X. doi: 10.7554/eLife.47088. URL https://doi.org/10.7554/eLife.47088.

Jorge Moura de Sousa, Roberto BalbontÃn, Paulo DurÃ£o, and Isabel Gordo. Multidrug-resistant bacteria compensate for the epistasis between resistances. PLOS Biology, 15(4):1–24, 04 2017. doi: 10.1371/journal.pbio.2001741. URL https://doi.org/10.1371/journal.pbio.2001741.

Philippe Remigi, Catherine Masson-Boivin, and Eduardo P.C. Rocha. Experimental evolution as a tool to investigate natural processes and molecular functions. Trends in Microbiology, 27(7):623–634, 2019. ISSN 0966-842X. doi: https://doi.org/10.1016/j.tim.2019.02.003. URL http://www.sciencedirect.com/science/article/pii/S0966842X19300411.

Rob J. Kulathinal, Brian R. Bettencourt, and Daniel L. Hartl. Compensated deleterious mutations in insect genomes. Science, 306(5701):1553–1554, 2004.

Robert Piskol and Wolfgang Stephan. Analyzing the evolution of RNA secondary structures in vertebrate introns using Kimura’s model of compensatory fitness interactions. Molecular Biology and Evolution, 25 (11):2483–2492, 2008.

Arthur Covert, Richard Lenski, Claus Wilke, and Charles Ofria. Experiments on the role of deleterious mutations as stepping stones in adaptive evolution. Proceedings of the National Academy of Sciences of the United States of America, 110(34):E3171–E3178, 2013.

Motoo Kimura. The role of compensatory neutral mutations in molecular evolution. Journal of Genetics, 64(1):7–19, 1985.

F. B. Moore, D. E. Rozen, and R. E. Lenski. Pervasive compensatory adaptation in Escherichia coli. Proceedings of the Royal Society of London B: Biological Sciences, 267(1442):515–522, 2000.

B. R. Levin, V. Perrot, and N. Walker. Compensatory mutations, antibiotic resistance and the population genetics of adaptive evolution in bacteria. Genetics, 154(3):985–997, 2000.

S. S. Choi, W. M. Li, and B. T. Lahn. Robust signals of coevolution of interacting residues in mammalian proteomes identified by phylogeny-aided structural analysis. Nature Genetics, 37(12):1367–1371, 2005.

Margarita V. Meer, Alexey S. Kondrashov, Yael Artzy-Randrup, and Fyodor A. Kondrashov. Compensatory evolution in mitochondrial trnas navigates valleys of low fitness. Nature, 464(7286):279–282, 2010.

S. Wright. Evolution in mendelian populations. Genetics, 16(2):97–159, 1931a.

S. Wright. The genetical theory of natural selection: A review. Journal of Heredity, 21(8):349–356, 1931b.

W. Stephan. The rate of compensatory evolution. Genetics, 144(1):419–426, 1996.

J. Parsch, S. Tanda, and W. Stephan. Site-directed mutations reveal long-range compensatory interactions in the Adh gene of Drosophila melanogaster. Proceedings of the National Academy of Sciences of the United States of America, 94(3):928–933, 1997.

M. C. Whitlock and S. P. Otto. The panda and the phage: Compensatory mutations and the persistence of small populations. Trends in Ecology & Evolution, 14(8):295–296, 1999.

M. C. Whitlock, C. K. Griswold, and A. D. Peters. Compensating for the meltdown: The critical effective size of a population with deleterious and compensatory mutations. Annales Zoologici Fennici, 40(2):169–183, 2003.

L. Zhang and L. T. Watson. Analysis of the fitness effect of compensatory mutations. HFSP Journal, 3(1): 47–54, 2009.

Brian Charlesworth. Effective population size and patterns of molecular evolution and variation. Nature Reviews Genetics, 10(3):195–205, 2009.

Sophie Maisnier-Patin, Otto G. Berg, Lars Liljas, and Dan I. Andersson. Compensatory adaptation to the deleterious effect of antibiotic resistance in salmonella typhimurium. Molecular Microbiology, 46(2): 355–366, 2002. ISSN 1365-2958. doi: 10.1046/j.1365-2958.2002.03173.x.

Danna R. Gifford and R. Craig MacLean. Evolutionary reversals of antibiotic resistance in experimental populations of Pseudomonas aeruginosa. Evolution, 67(10):2973–2981, 2013. ISSN 1558-5646. doi: 10.1111/evo.12158.

Daniel B Sloan, Deborah A Triant, Martin Wu, and Douglas R Taylor. Cytonuclear interactions and relaxed selection accelerate sequence evolution in organelle ribosomes. Molecular Biology and Evolution, 31(3): 673–682, March 2014.

Ricardo B. R. Azevedo, Rolf Lohaus, Suraj Srinivasan, Kristen K. Dang, and Christina L. Burch. Sexual reproduction selects for robustness and negative epistasis in artificial gene networks. Nature, 440(7080): 87–90, 2006.

A. Wagner. Does evolutionary plasticity evolve? Evolution, 50(3):1008–1023, 1996.

M. L. Siegal and A. Bergman. Waddington’s canalization revisited: Developmental stability and evolution. Proceedings of the National Academy of Sciences of the United States of America, 99(16):10528–10532, 2002.

R Lohaus, C L Burch, and R B R Azevedo. Genetic architecture and the evolution of sex. The Journal of Heredity, 101(Supplement 1):S142–S157, April 2010.

Yifei Wang, Yinghong Lan, Daniel M. Weinreich, Nicholas K. Priest, and Joanna J. Bryson. Recombination is surprisingly constructive for artificial gene regulatory networks in the context of selection for developmental stability. In Proceedings of the European Conference on Artificial Life 2015, pages 530–537. The MIT Press, July 2015.

Yifei Wang. Review of wagner’s artificial gene regulatory networks model and its applications for understanding complex biological systems. COJ Robotics & Artificial Intelligence, 1(1):COJRA.000501.2019, Feb 2019a.

Yifei Wang. Convergence analysis and network properties of wagner’s artificial gene regulatory network model. Journal of Advances in Mathematics and Computer Science, 31(2):1–18, Mar 2019b.

H. Crawford, J. G. Prado, A. Leslie, S. Hue, I. Honeyborne, S. Reddy, M. van der Stok, Z. Mncube, C. Brander, C. Rousseau, J. I. Mullins, R. Kaslow, P. Goepfert, S. Allen, E. Hunter, J. Mulenga, P. Kiepiela, B. D. Walker, and P. J. Goulder. Compensatory mutation partially restores fitness and delays reversion of escape mutation within the immunodominant HLA-B*5703-restricted Gag epitope in chronic human immunodeficiency virus type 1 infection. Journal of Virology, 81(15):8346–8351, 2007.

Tiffany B. Taylor, Geraldine Mulley, Alexander H. Dills, Abdullah S. Alsohim, Liam J. McGuffin, David J. Studholme, Mark W. Silby, Michael A. Brockhurst, Louise J. Johnson, and Robert W. Jackson. Evolutionary resurrection of flagellar motility via rewiring of the nitrogen regulation system. Science, 347(6225): 1014–1017, 2015. ISSN 0036-8075. doi: 10.1126/science.1259145.

Daniel M. Weinreich and Lin Chao. Rapid evolutionary escape by large populations from local fitness peaks is likely in nature. Evolution, 59(6):1175–1182, 2005. doi: 10.1111/j.0014-3820.2005.tb01769.x. URL https://onlinelibrary.wiley.com/doi/abs/10.1111/j.0014-3820.2005.tb01769.x.

Inaki Comas, Sonia Borrell, Andreas Roetzer, Graham Rose, Bijaya Malla, Midori Kato-Maeda, James Galagan, Stefan Niemann, and Sebastien Gagneux. Whole-genome sequencing of rifampicin-resistant mycobacterium tuberculosis strains identifies compensatory mutations in RNA polymerase genes. Nature Genetics, 44(1):106–110, 2012.

Gerrit Brandis, Marie Wrande, Lars Liljas, and Diarmaid Hughes. Fitness-compensatory mutations in rifampicin-resistant RNA polymerase. Molecular Microbiology, 85(1):142–151, July 2012.

M de Vos, B Mueller, S Borrell, P A Black, P D van Helden, R M Warren, S Gagneux, and T C Victor. Putative compensatory mutations in the rpoC gene of rifampin-resistant Mycobacterium tuberculosis are associated with ongoing transmission. Antimicrobial Agents and Chemotherapy, 57(2):827–832, February 2013.

Gerrit Brandis and Diarmaid Hughes. Genetic characterization of compensatory evolution in strains carrying rpoB Ser531Leu, the rifampicin resistance mutation most frequently found in clinical isolates. Journal of Antimicrobial Chemotherapy, 68(11):2493–2497, November 2013.

Taeksun Song, Yumi Park, Isdore Chola Shamputa, Sunghwa Seo, Sun Young Lee, Han-Seung Jeon, Hongjo Choi, Myungsun Lee, Richard J Glynne, S Whitney Barnes, John R Walker, Serge Batalov, Karina Yusim, Shihai Feng, Chang-Shung Tung, James Theiler, Laura E Via, Helena I M Boshoff, Katsuhiko S Murakami, Bette Korber, Clifton E III Barry, and Sang-Nae Cho. Fitness costs of rifampicin resistance in Mycobacterium tuberculosis are amplified under conditions of nutrient starvation and compensated by mutation in the *β^l^* subunit of RNA polymerase. Molecular Microbiology, 91(6):1106–1119, March 2014.

Carlos Martinez, Joshua S Rest, Ah-Ram Kim, Michael Ludwig, Martin Kreitman, Kevin White, and John Reinitz. Ancestral resurrection of the Drosophila S2E enhancer reveals accessible evolutionary paths through compensatory change. Molecular Biology and Evolution, 31(4):903–916, April 2014.

Dmitry N Ivankov, Alexei V Finkelstein, and Fyodor A Kondrashov. A structural perspective of compensatory evolution. Current Opinion in Structural Biology, 26:104–112, June 2014.

Bela Szamecz, Gabor Boross, Dorottya Kalapis, Karoly Kovacs, Gergely Fekete, Zoltan Farkas, Viktoria Lazar, Monika Hrtyan, Patrick Kemmeren, Marian J A Groot Koerkamp, Edit Rutkai, Frank C P Holstege, Balazs Papp, and Csaba Pal. The genomic landscape of compensatory evolution. PLoS Biology, 12(8): e1001935, August 2014.

Marie Filteau, Véronique Hamel, Marie-Christine Pouliot, Isabelle Gagnon-Arsenault, Alexandre K Dubé, and Christian R Landry. Evolutionary rescue by compensatory mutations is constrained by genomic and environmental backgrounds. Molecular Systems Biology, 11(10), 2015. ISSN 1744-4292. doi: 10.15252/msb.20156444.

A San Millan, R Peña-Miller, M Toll-Riera, Z V Halbert, A R McLean, B S Cooper, and R C MacLean. Positive selection and compensatory adaptation interact to stabilize non-transmissible plasmids. Nature Communications, 5:5208, 2014.

Joanne L Porter, Charles A Collyer, and David L Ollis. Compensatory stabilizing role of surface mutations during the directed evolution of dienelactone hydrolase for enhanced activity. Protein Journal, 34(1): 82–89, February 2015.

Ellie Harrison, David Guymer, Andrew J Spiers, Steve Paterson, and Michael A Brockhurst. Parallel compensatory evolution stabilizes plasmids across the parasitism-mutualism continuum. Current Biology, 25 (15):1–6, July 2015.

Naoki Osada and Hiroshi Akashi. Mitochondrial-nuclear interactions and accelerated compensatory evolution: Evidence from the primate cytochrome C oxidase complex. Molecular Biology and Evolution, 29(1): 337–346, January 2012.

Felipe S Barreto and Ronald S Burton. Evidence for compensatory evolution of ribosomal proteins in response to rapid divergence of mitochondrial rRNA. Molecular Biology and Evolution, 30(2):310–314, February 2013.

S. A. West, A. D. Peters, and N. H. Barton. Testing for epistasis between deleterious mutations. Genetics, 149(1):435–444, 1998.

West, Lively, and Read. A pluralist approach to sex and recombination. Journal of Evolutionary Biology, 12(6):1003–1012, 1999.

Joanna J. Bryson, Yasushi Ando, and Hagen Lehmann. Agent-based models as scientific methodology: A case study analysing primate social behaviour. Philosophical Transactions of the Royal Society of London B: Biological Sciences, 362(1485):1685–1698, September 2007.

Adam M. Siepielski, Joseph D. DiBattista, and Stephanie M. Carlson. It’s about time: The temporal dynamics of phenotypic selection in the wild. Ecology Letters, 12(11):1261–1276, 2009b. ISSN 1461-0248. doi: 10.1111/j.1461-0248.2009.01381.x.

Benjamin Brachi, Romain Villoutreix, Nathalie Faure, Nina Hautekèete, Yves Piquot, Maxime Pauwels, Dominique Roby, Joël Cuguen, Joy Bergelson, and Fabrice Roux. Investigation of the geographical scale of adaptive phenological variation and its underlying genetics in Arabidopsis thaliana. Molecular Ecology, 22(16):4222–4240, 2013. ISSN 1365-294X. doi: 10.1111/mec.12396.

Zachariah Gompert, Aaron A. Comeault, Timothy E. Farkas, Jeffrey L. Feder, Thomas L. Parchman, C. Alex Buerkle, and Patrik Nosil. Experimental evidence for ecological selection on genome variation in the wild. Ecology Letters, 17(3):369–379, 2014. ISSN 1461-0248. doi: 10.1111/ele.12238.

O. Seppälä. Natural selection on quantitative immune defence traits: A comparison between theory and data. Journal of Evolutionary Biology, 28(1):1–9, 2015. ISSN 1420-9101. doi: 10.1111/jeb.12528.

Allert I. Bijleveld, Sönke Twietmeyer, Julia Piechocki, Jan A. van Gils, and Theunis Piersma. Natural selection by pulsed predation: Survival of the thickest. Ecology, 96(7):1943–1956, 2015.

Caitlin F Connelly, Jon Wakefield, and Joshua M Akey. Evolution and genetic architecture of chromatin accessibility and function in yeast. PLoS Genetics, 10(7):e1004427, July 2014.

C Espinosa-Soto, O C Martin, and A Wagner. Phenotypic robustness can increase phenotypic variability after nongenetic perturbations in gene regulatory circuits. Journal of Evolutionary Biology, 24(6):1284–1297, June 2011.

Rafael Sanjuán, Andrés Moya, and Santiago F. Elena. The distribution of fitness effects caused by single-nucleotide substitutions in an RNA virus. Proceedings of the National Academy of Sciences of the United States of America, 101(22):8396–8401, 2004.

Adam Eyre-Walker and Peter Keightley. The distribution of fitness effects of new mutations. Nature Reviews Genetics, 8(8):610–618, 2007.

Peter D. Keightley and Adam Eyre-Walker. Joint inference of the distribution of fitness effects of deleterious mutations and population demography based on nucleotide polymorphism frequencies. Genetics, 177(4): 2251–2261, 2007.

S. Mezmouk and J. Ross-Ibarra. The pattern and distribution of deleterious mutations in maize. G3 (Bethesda, Md.), 4(1):163–171, 2014.

A. Poon, B. H. Davis, and L. Chao. The coupon collector and the suppressor mutation: Estimating the number of compensatory mutations by maximum likelihood. Genetics, 170(3):1323–1332, 2005.

A. Poon and L. Chao. The rate of compensatory mutation in the DNA bacteriophage phi X174. Genetics, 170(3):989–999, 2005.

Brad H. Davis, Art F. Y. Poon, and Michael C. Whitlock. Compensatory mutations are repeatable and clustered within proteins. Proceedings of the Royal Society of London B: Biological Sciences, 276(1663): 1823–1827, 2009.

Amrita Bhattacherjee, Saurav Mallik, and Sudip Kundu. Compensatory mutations occur within the electrostatic interaction range of deleterious mutations in protein structure. Journal of Molecular Evolution, 80(1):10–12, January 2015.

Troy Ruths and Luay Nakhleh. Neutral forces acting on intragenomic variability shape the Escherichia coli regulatory network topology. Proceedings of the National Academy of Sciences of the United States of America, 110(19):7754–7759, 2013.

Joshua L. Payne and Andreas Wagner. Mechanisms of mutational robustness in transcriptional regulation. Frontiers in Genetics, 6(322), 2015.

Andreas Wagner. Evolution of gene networks by gene duplications: A mathematical model and its implications on genome organization. Proceedings of the National Academy of Sciences of the United States of America, 91(10):4387–4391, 1994.

